# Evolutionary persistence and divergence of the *tdk* killer meiotic driver family

**DOI:** 10.64898/2025.12.28.696746

**Authors:** Fan-Yi Zhang, Guo-Song Jia, Jing-Yi Ren, Fang Suo, Tong-Yang Du, Wen-Cai Zhang, Wen Li, Qi-Ming Wang, Li-Lin Du, Yu Hua

## Abstract

Killer meiotic drivers (KMDs) are selfish genetic elements that achieve super-Mendelian inheritance by selectively eliminating gametes lacking the driver. Although predicted to arise recurrently, KMDs are generally considered evolutionarily ephemeral—going extinct after fixation or host suppression. The identification of *tdk1*, a single-gene KMD in the fission yeast *Schizosaccharomyces pombe*, provides a model for studying KMD evolution. Here, we identify two divergent *tdk1* homologs (*tdk210* and *tdk203*) from *S. cryophilus*, a fission yeast species that diverged ∼100 million years ago from *S. pombe*, as active KMDs. These three KMDs all act via post-germination killing, disrupting chromosome segregation in noncarrier progeny. Notably, they also exhibit striking functional divergences: *tdk1*, *tdk210*, and *tdk203* are mutually incompatible (showing no cross-resistance), and the latter two act independently of Bdf1/Bdf2—host chromatin proteins required for *tdk1* killing. Phylogenetic analyses of the dozens of *tdk* family genes in *Schizosaccharomyces* support long-term persistence and rapid evolutionary dynamics of this gene family. Remarkably, homologs in distantly related fungal phyla display genomic and structural similarities to *Schizosaccharomyces tdk* genes, suggesting a deeply rooted origin of this KMD family in fungi. Our findings reveal that a single KMD family can undergo repeated functional innovation—generating mutually incompatible variants and rewiring host dependencies—while maintaining a conserved killing mode over deep evolutionary time.

## INTRODUCTION

Genomes are battlegrounds for selfish genetic elements that prioritize their own transmission, often at the expense of host fitness^1–5^. Among these, killer meiotic drivers (KMDs) act with striking ruthlessness: in driver+/driver− heterozygotes, they kill or disable gametes lacking the driver allele, thereby skewing Mendelian transmission in their favor^6^. This transmission advantage allows KMDs to spread rapidly in sexual populations—even as they impose fitness costs on hosts—fueling genetic conflicts that shape genome architecture, speciation, and the evolution of sexual reproduction^1,7–10^.

A long-standing view holds that while KMDs arise recurrently in eukaryotes, individual drive systems are evolutionarily ephemeral and restricted to narrow phylogenetic ranges^1,6,7,11,12^. This transience is thought to stem from the fragility of their selective advantage^1,6,7^. Specifically, the fitness costs of KMDs select for host suppressors that counteract drive; alternatively, if a KMD reaches fixation before suppressor emergence, the heterozygous context required for drive is lost. In both cases, deprived of their selective advantage, KMDs degenerate and go extinct. Recent findings, however, have begun to challenge this long-held view by uncovering KMD families with broad phylogenetic distributions^13^. Notable examples include the *Spok* family (first discovered in *Podospora anserina*), whose homologs span multiple filamentous fungal lineages^14–16^, and the *wtf* family in fission yeasts—estimated to have persisted for over 100 million years^17–19^.

The recent identification of *tdk1*, a single-gene KMD in the fission yeast *Schizosaccharomyces pombe*, provides a new model for investigating KMD evolution. The *tdk1* gene encodes a protein that adopts dual conformations, functioning as a toxin–antidote system^20,21^. The toxin conformation is present in all meiotic products (i.e., spores in fungi)^21^, whereas the antidote—which neutralizes the toxin—is produced exclusively in progeny carrying *tdk1*, leading to selective elimination of noncarrier progeny^20^. Intriguingly, distant Tdk1 homologs (<30% amino acid identity) have been identified across evolutionarily divergent *Schizosaccharomyces* species^21^. However, it remains unclear whether these homologs represent non-driver precursors of *tdk1* or, alternatively, members of an ancient, highly diverged KMD family. If the latter, what evolutionary mechanisms have enabled their long-term persistence?

To address these questions, we performed functional and phylogenetic analyses of *tdk1* homologs. We identified two divergent homologs from *S. cryophilus*—a fission yeast species that diverged ∼100 million years ago (Mya) from *S. pombe*^22^—as active KMDs, demonstrating that *tdk1* and its fission yeast homologs constitute a KMD family—designated the *tdk* gene family. By dissecting the conserved and divergent features of the three active *tdk* drivers, we uncover molecular strategies that enable their persistence over deep evolutionary time. In addition, leveraging recently generated telomere-to-telomere (T2T) assemblies of nine representative *Schizosaccharomyces* genomes (Jia et al., manuscript in preparation), we comprehensively characterized the distribution and evolutionary dynamics of *tdk* family genes within this genus. Moreover, we identified *tdk* homologs in disparate fungal phyla that share genomic and structural features with *Schizosaccharomyces tdk* genes, supporting a deep evolutionary origin of this KMD family in fungi.

## RESULTS

### Identification of *S. cryophilus tdk210* and *tdk203* as active killer meiotic drivers

We previously identified and characterized *tdk1* in *S. pombe* as a single-gene KMD, whose protein product adopts two antagonistic conformations to function as a toxin–antidote system^20,21^. The toxin conformation is formed in all meiotic haploid progeny and disrupts post-germination mitosis by causing aberrant chromosomal adhesions via interactions with the host chromatin proteins Bdf1 and Bdf2^21^. In contrast, the antidote — corresponding to newly synthesized Tdk1 protein in *tdk1*-carrying progeny—neutralizes the toxic form^20^.

Our prior BLAST search of Tdk1 against the NCBI RefSeq database identified 15 homologous proteins across divergent fission yeast species (*S. pombe*, *S. cryophilus*, *S. osmophilus*, and *S. octosporus*)^21^. Given the low sequence identity between Tdk1 and these homologs (18.2%–28.7% amino acid identity; Supplementary Fig. 1a) and the typical confinement of KMDs to single species or narrow phylogenetic lineages, a critical question arises: is *tdk1* a recently evolved driver derived from non-driver precursors in *S. pombe*, or a member of a highly diverged ancient KMD family?

To address this, we functionally tested all 15 homologs of *tdk1* for meiotic drive activity. We conducted the analysis in *S. pombe* owing to its genetic tractability, although heterologous expression may fail to recapitulate drive activity if it requires species-specific context. Each homolog’s coding sequence was cloned under the control of the *tdk1* promoter and terminator (*P.tdk1* and *T.tdk1*) and integrated at the *ade6* locus. Haploid strains carrying these constructs were crossed to a strain lacking the insertion. Of the 15 homologs tested, two from *S. cryophilus*—*SPOG_02247* and *SPOG_05700* (27.7% and 27.3% amino acid identity to Tdk1, respectively), designated *tdk210* and *tdk203* (see below)—exhibited substantial killing of noncarrier progeny in heterozygous crosses. Noncarrier viability was reduced to 4.2% for *tdk210* and 45.2% for *tdk203*, whereas carrier progeny exhibited normal viability (∼90%) (Supplementary Fig. 1b). *tdk210* and *tdk203* remained active in *S. pombe* when expressed under their native *S. cryophilus* regulatory sequences (promoters and terminators) — in heterozygous crosses, noncarrier progeny showed low viability (14.2% for *tdk210*; 20.4% for *tdk203*), whereas carrier progeny exhibited normal viability (Fig. 1a,b). As controls, homozygous crosses for *tdk210* or *tdk203* consistently exhibited normal progeny viability (Fig. 1a,b). These results demonstrate that *tdk210* and *tdk203* from *S. cryophilus* are active KMDs and support the notion that *tdk1* and its homologs constitute a KMD family, which we designate the *tdk* family.

**Fig. 1:**
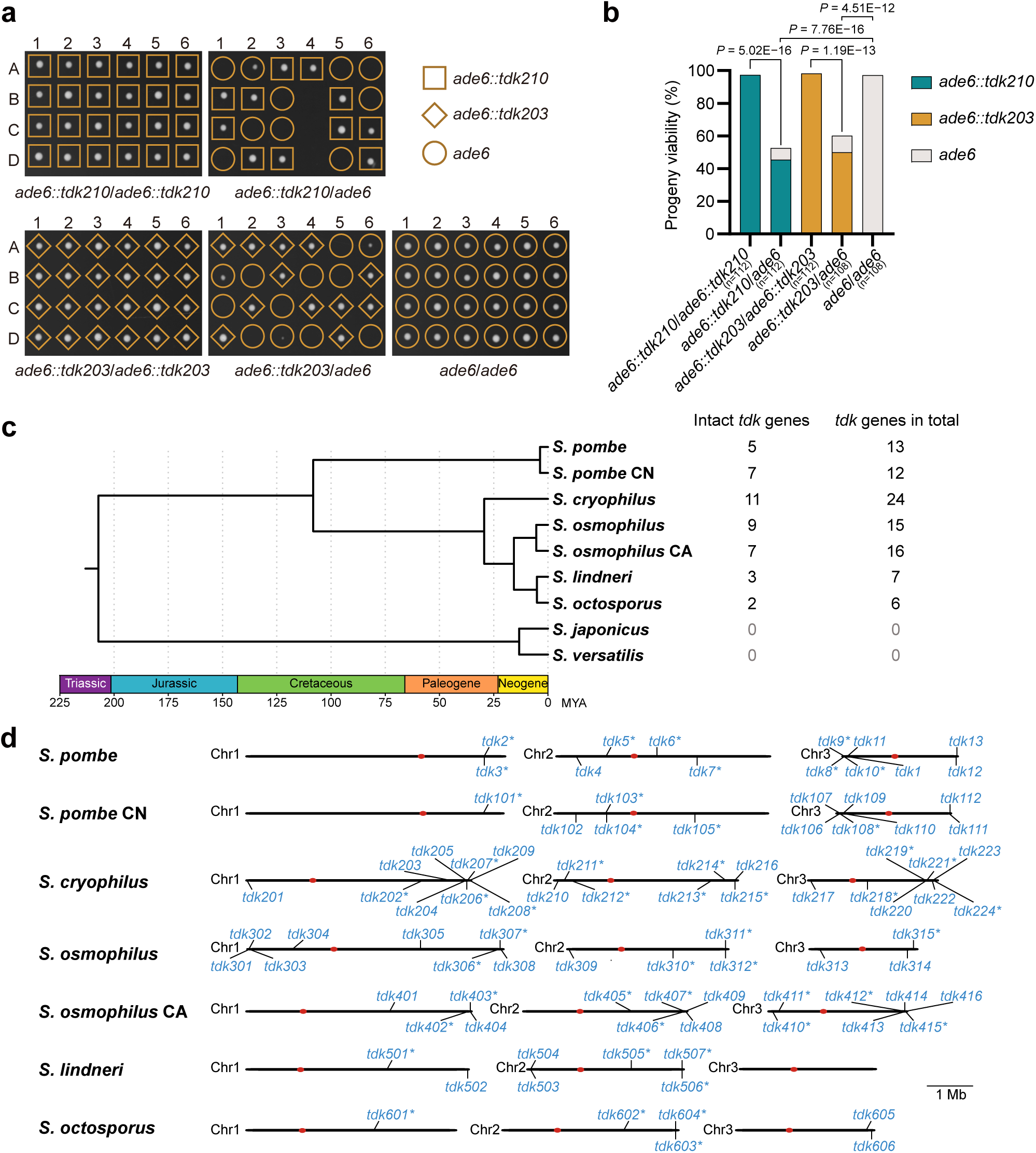
Identification of *S. cryophilus tdk210* and *tdk203* as active KMDs among *Schizosaccharomyces tdk* homologs. **a,b**, Tetrad analyses of *ade6::tdk210* × *ade6::tdk210*, *ade6::tdk210* × *ade6*, *ade6::tdk203* × *ade6::tdk203*, *ade6::tdk203* × *ade6*, and *ade6* × *ade6* (control) crosses showing inviability of noncarrier progeny from heterozygous diploids. **a**, Six representative tetrads per cross; A–D denote progeny from a single tetrad. **b**, Statistical analysis of progeny viability. *P* values (Fisher’s exact test) compare progeny viability between crosses (null hypothesis: equal viability). n, total progeny analyzed. *tdk210* and *tdk203* genes (with native promoters and terminators) were inserted at the *ade6* locus in the *tdk1Δ tdk11Δ* background. **c**, Time-calibrated phylogeny of the *Schizosaccharomyces* genus (Jia et al., manuscript in preparation), with numbers of intact *tdk* genes and total *tdk* genes (including pseudogenes) listed on the right. **d**, Genomic maps of *tdk* loci in *S. pombe*, *S. pombe* CN, *S. cryophilus*, *S. osmophilus*, *S. osmophilus* CA, *S. lindneri*, and *S. octosporus*. Predicted pseudogenes are marked with an asterisk (*). Full locus information is provided in Supplementary Data 1.

### Systematic identification and annotation of *tdk* family genes in *Schizosaccharomyces*

To systematically characterize the *tdk* gene repertoire across *Schizosaccharomyces*, we analyzed T2T assemblies of nine representative fission yeast genomes (Jia et al., manuscript in preparation). These include the genomes of the reference strains of seven fission yeast species: *S. pombe*, *S. cryophilus*, *S. osmophilus*, *S. lindneri*, and *S. octosporus* — which share a common ancestor ∼100 Mya — along with two more divergent species, *S. japonicus* and *S. versatilis*, that separated from the others ∼200 Mya^22^. Two genomes representing highly diverged intraspecific lineages (*S. pombe* CN and *S. osmophilus* CA) were also included. The sequence of *tdk203*—incompletely assembled in RefSeq—was corrected using the new genome assembly of *S. cryophilus*.

We searched for *tdk* family genes using PSI-BLAST^23^ against the predicted proteins encoded by the nine genomes. Searches with Tdk1, Tdk210, or Tdk203 as queries yielded identical candidate sets, including the 15 *tdk1* homologs identified in the prior BLAST search against RefSeq. *S. japonicus* and *S. versatilis* lacked detectable PSI-BLAST hits; all other genomes yielded multiple hits. By performing sequence alignments of the PSI-BLAST hits, we noticed that a substantial portion of them appeared to be pseudogenes that do not encode intact proteins. Thus, we conducted manual curation to distinguish intact genes from pseudogenes using long-read transcriptomic data and sequence alignments (see Methods).

In *S. pombe*, PomBase currently annotates eight *tdk* family members in addition to *tdk1*: *meu23*, *mug2*, *new22*, *SPBC337.02c*, *SPCP20C8.01c*, *SPCC569.01c*, *SPCC569.03*, and one pseudogene (*SPBC18E5.15*). Our manual curation led to three revisions: *SPBC337.02c*, *SPCP20C8.01c*, and *new22* were reclassified as pseudogenes. One *S. cryophilus* gene (*SPOG_03615*), which is among the 15 previously identified *tdk1* homologs, was also reclassified as a pseudogene. A total of 44 intact *tdk* genes remain after the manual curation: 5 in *S. pombe*, 7 in *S. pombe* CN, 11 in *S. cryophilus*, 9 in *S. osmophilus*, 7 in *S. osmophilus* CA, 3 in *S. lindneri*, and 2 in *S. octosporus* (Fig. 1c,d and Supplementary Data 1).

To more comprehensively detect pseudogenic *tdk* loci, we performed tBLASTn searches using the protein sequences encoded by the 44 intact *tdk* family genes as queries. Again, no homologous sequences were found in *S. japonicus* or *S. versatilis*. Four additional *tdk* pseudogenes were identified in *S. pombe*: *SPAC29A4.23*, *SPCP20C8.03*, and two unannotated genes (we assigned them systematic IDs *SPA2025.04* and *SPB2025.27*). In total, we identified 93 *tdk* family members (including 44 intact genes and 49 pseudogenes) across the seven other genomes: 13 in *S. pombe*, 12 in *S. pombe* CN, 24 in *S. cryophilus*, 15 in *S. osmophilus*, 16 in *S. osmophilus* CA, 7 in *S. lindneri*, and 6 in *S. octosporus* (Fig. 1c,d and Supplementary Data 1). Notably, *tdk* loci are enriched near chromosome ends: 57 of the 93 *tdk* genes are located within the terminal 5% of chromosomal contigs (Fig. 1d).

To facilitate future research, we established a standardized nomenclature for *tdk* family genes across *Schizosaccharomyces*. In *S. pombe*, all *tdk* loci (except *tdk1*) were named according to chromosomal position: on Chr I, *SPAC29A4.23* and *SPA2025.04* were designated *tdk2\** and *tdk3*\* (* denotes pseudogenes), respectively; on Chr II, *mug2* was renamed *tdk4*, and *SPBC337.02c*, *SPBC18E5.15*, and *SPB2025.27* were assigned *tdk5\**, *tdk6\**, and *tdk7\**; on Chr III, *SPCP20C8.01c*, *SPCP20C8.03*, *new22*, *meu23*, *SPCC569.03*, and *SPCC569.01c* were designated *tdk8\**, *tdk9\**, *tdk10\**, *tdk11*, *tdk12*, and *tdk13*, respectively (Fig. 1d and Supplementary Data 1).

For the remaining six *Schizosaccharomyces* genomes, we assigned each *tdk* gene a unique three-digit identifier. Due to limited synteny conservation (see below), homologs within each genome were named in sequential order along the chromosomes, proceeding from the left arm of Chr I to the right arm of Chr III. The first digit indicates the species or lineage (1 = *S. pombe* CN; 2 = *S. cryophilus*; 3 = *S. osmophilus*; 4 = *S. osmophilus* CA; 5 = *S. lindneri*; 6 = *S. octosporus*), while the final two digits reflect the gene’s sequential position along the chromosomes (Fig. 1d). The two active KMDs in *S. cryophilus* were thus designated *tdk210* and *tdk203*, respectively.

The more comprehensive gene inventory of the *tdk* family enabled us to test six additional members—*tdk222*, *tdk223*, *tdk302*, *tdk305*, *tdk309*, and *tdk313*—from *S. cryophilus* and *S. osmophilus*, the two species with the most intact *tdk* repertoires; however, none exhibited drive activity (Supplementary Fig. 1c).

### *tdk1*, *tdk210*, and *tdk203* are active when expressed from a strong constitutive promoter

To determine whether *tdk210* and *tdk203* require precise expression control for activity, we first verified the correctness of their annotated coding sequences. Start-codon mutations (*tdk210-M1A* and *tdk203-M1A*) abolished killing in crosses with noncarriers (Fig. 2a), confirming correct annotation. In crosses with wild-type carriers, *ade6::tdk210-M1A* progeny showed low viability (18.2%), comparable to noncarriers, indicating loss of protection. In contrast, *ade6::tdk203-M1A* progeny exhibited unexpectedly high viability (87.6%), possibly due to downstream translation initiation producing a protective truncated protein.

**Fig. 2:**
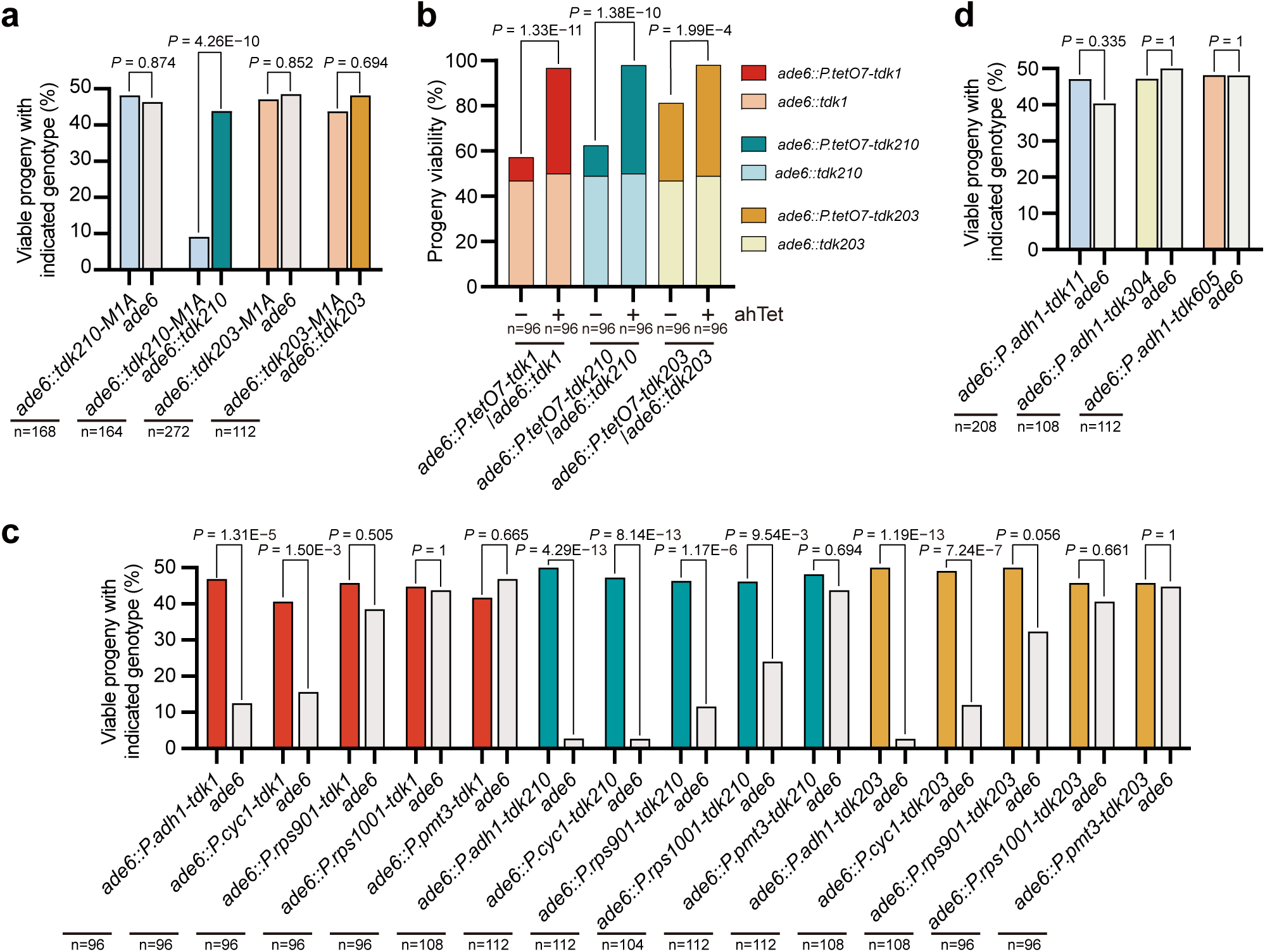
*tdk* drivers share a conserved single-protein toxin–antidote mechanism. **a**, Tetrad analyses of *ade6::tdk210-M1A* × *ade6*, *ade6::tdk210-M1A* × *ade6::tdk210*, *ade6::tdk203-M1A* × *ade6*, and *ade6::tdk203-M1A* × *ade6::tdk203* crosses, showing that the start codon mutation (M1A) abolishes killing activity of *tdk210* and *tdk203* and the antidote (resistance) activity of *tdk210*; the unexpected resistance of *tdk203-M1A* against *tdk203* may arise from downstream translation initiation. **b**, Tetrad analyses of *ade6::P.tetO7-tdk1* × *ade6::tdk1*, *ade6::P.tetO7-tdk210* × *ade6::tdk210* and *ade6::P.tetO7-tdk203* × *ade6::tdk203* crosses, showing that inducing *tdk* expression during germination via the *P.tetO7* promoter (by transferring spores to a germination medium containing anhydrotetracycline [ahTet]) rescued the progeny killing by *ade6::tdk*. **c**, Tetrad analyses demonstrating dosage-dependent killing of noncarrier progeny by *tdk1*, *tdk210*, and *tdk203* expressed from constitutive promoters of decreasing strength (*P.adh1*, *P.cyc1*, *P.rps901*, *P.rps1001*, and *P.pmt3*). Promoter strength was validated as shown in Supplementary Fig. 2c. **d**, Tetrad analyses of *ade6::P.adh1-tdk11* × *ade6*, *ade6::P.adh1-tdk304* × *ade6*, and *ade6::P.adh1-tdk605* × *ade6* crosses, showing marginal drive activity for *P.adh1-tdk11* but no activity for *P.adh1-tdk304* or *P.adh1-tdk605*. For **a,c,d**, *P* values (exact binomial test) compare observed viable progeny counts of the two genotypes to the expected 1:1 Mendelian segregation ratio. For **b**, *P* values (Fisher’s exact test) compare progeny viability between crosses. n, total progeny analyzed.

We next asked whether driver expression during spore germination is sufficient for protection—a property established for *tdk1*^20^. Using an anhydrotetracycline (ahTet)-inducible promoter (*P.tetO7*^24^), we found that ahTet-induced expression of *tdk1*, *tdk210*, or *tdk203* during germination fully restored progeny viability (>90%), whereas uninduced expression resulted in low viability (20.8%, 27.0%, and 68.8%, respectively; Fig. 2b). The relatively high baseline viability for *tdk203* likely reflects promoter leakiness.

Because *tdk1* is meiotically upregulated via a Mei4-binding motif^20^, we asked whether *tdk210* and *tdk203* are similarly regulated. However, neither *tdk210* nor *tdk203* promoters contain Mei4 sites, and neither shows meiotic upregulation (Supplementary Fig. 2a), suggesting such regulation is dispensable. To test this directly, we expressed all three drivers under the strong constitutive *P.adh1* promoter^25^. *P.adh1*-driven expression caused no vegetative toxicity (Supplementary Fig. 2b) but drastically reduced noncarrier viability in crosses (25.0% for *tdk1*; 5.6% for *tdk210*; 5.4% for *tdk203*; Fig. 2c), demonstrating robust drive. Further experiments with constitutive promoters of decreasing strength revealed a dosage effect: weaker promoters generally exhibited reduced drive activity, though the minimal promoter strength sufficient for robust drive differed among the three *tdks* (Fig. 2c and Supplementary Fig. 2c).

Given that *tdk210* and *tdk203* exhibit stronger drive activity under *P.adh1* than their native promoters, we tested whether their closest homologs, which did not exhibit obvious drive activities when expressed under their native promoters, are capable of drive under *P.adh1*. These homologs included: *tdk11* (51.4% amino acid identity to Tdk210) and two close relatives of *tdk203*—*tdk304* (68.5% identity to Tdk203) and *tdk605* (69.3% identity to Tdk203). Only *tdk11* showed marginal drive (noncarrier viability: 80.8% vs. 94.2% for carriers); *tdk304* and *tdk605* remained inactive (Fig. 2d). Together, these results indicate that active *tdk* drivers do not require elaborate transcriptional regulation, as constitutive expression is sufficient for drive. However, the inability of *tdk304* and *tdk605* to drive — even under strong constitutive expression—suggests that their inactivity stems from intrinsic functional defects, not merely low expression.

### Tdk proteins share conserved structural features

To assess structural conservation among the protein products of the three active drivers (*tdk1*, *tdk210*, and *tdk203*) as well as the marginal driver *tdk11*, we generated AlphaFold 3 predictions^26^. The predicted aligned error (PAE) plots provided by AlphaFold 3 indicated that the four proteins all adopt a bipartite architecture comprising two independently folded regions connected by a short flexible linker, with a conserved PP motif positioned at the linker (Fig. 3a and Supplementary Fig. 3a–d). The predicted structures of the N-terminal regions all feature a portion that adopts a distinctive Z-shape (residues 12–45 in Tdk1, hereafter referred to as the Z-shaped domain) (Fig. 3b–d and Supplementary Fig. 3e). The C-terminal regions all contain a conserved InterPro domain (IPR013902; Pfam PF08593), corresponding to residues 269–340 in Tdk1—the most sequence-conserved segment—and an α-helix that immediately precedes this domain, whose length varies among Tdk proteins (Fig. 3e–g and Supplementary Fig. 3 a–d,f).

**Fig. 3:**
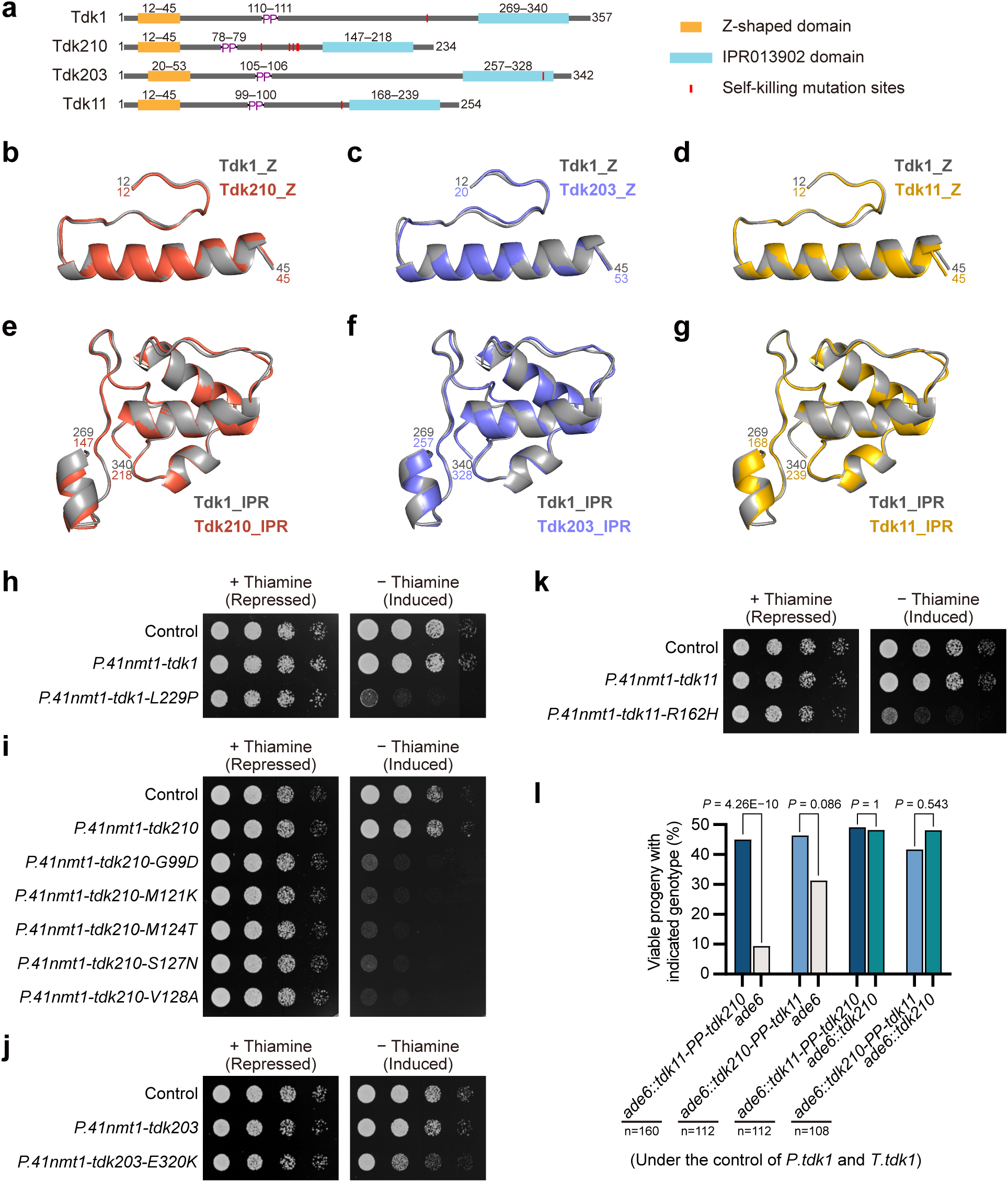
Tdk driver proteins share conserved structural features. **a**, Schematic of Tdk1, Tdk210, Tdk203, and Tdk11, highlighting the conserved Z-shaped domain, PP motif (connecting N- and C-terminal regions), IPR013902 domain, and point mutation sites, each of which confers self-killing activity, as shown in Fig. 2h–k. **b–g**, Structural alignments of the Z-shaped domain (**b–d**) and IPR013902 domain (**e–g**) of Tdk210 (**b,e**; Z-domain RMSD = 0.314 Å over 228 atoms; IPR013902 domain RMSD = 0.324 Å over 361 atoms), Tdk203 (**c,f**; Z-domain RMSD = 0.296 Å over 168 atoms; IPR013902 domain RMSD = 0.374 Å over 404 atoms), and Tdk11 (**d,g**; Z-domain RMSD = 0.289 Å over 186 atoms; IPR013902 domain RMSD = 0.245 Å over 386 atoms) to Tdk1 (dark gray). Structures were predicted by AlphaFold 3^26^; full models and predicted aligned error (PAE) plots are provided in Supplementary Fig. 3a–d. **h–k**, Spot assays showing vegetative toxicity of self-killing *tdk* mutants (*tdk1-L229P*; *tdk210-G99D*/*M121K*/*M124K*/*S127N*/*V128A*; *tdk203-E320K*; *tdk11-R162H*) but not their wild-type counterparts, under control of the *P.nmt41* promoter. **l**, Tetrad analyses of crosses involving *tdk11***–***tdk210* chimeras (*tdk11-PP-tdk210* and *tdk210-PP-tdk11*,each controlled by *P.tdk1* and *P.tdk1*), demonstrating that both chimeras retain drive activity against noncarriers and confer resistance to *tdk210*. *P* values (exact binomial test) compare progeny viability of the two genotypes within each cross. n, total progeny analyzed.

A single point mutation (L229P) converts Tdk1 into a self-killing variant in vegetative cells^21^. To determine whether Tdk210, Tdk203, and Tdk11 could similarly be converted into self-killing variants by point mutations — despite the lack of conservation of Tdk1 Leu229 among these proteins — we used rolling circle amplification mutagenesis^27,28^, and identified at least one such mutation in each *tdk* driver: G99D, M121K, M124K, S127N, or V128A in *tdk210*; E320K in *tdk203*; and R162H in *tdk11* (see Methods). When expressed from the *P.41nmt1* promoter, each mutant conferred vegetative toxicity (Fig. 3h–k). All but one of the self-killing mutations (E320K in *tdk203*) mapped to the α-helix that precedes the IPR013902 domain (Fig. 3a). These results suggest that all four Tdk proteins are prevented from adopting a toxic conformation in vegetative cells by a structural constraint that can be disrupted by a single point mutation.

To further probe structural conservation, we generated reciprocal chimeric proteins by swapping the N- and C-terminal regions at the conserved PP motif. Notably, chimeras between Tdk210 and its closest homolog Tdk11 (51.4% amino acid identity)—*tdk11-PP-tdk210* and *tdk210-PP-tdk11*—both exhibited substantial drive activity, reducing noncarrier viability to 18.8% and 62.6%, respectively, and conferring resistance to *tdk210* (Fig. 3l). In contrast, chimeras between more divergent pairs—Tdk11 and Tdk1 (28.7% identity) or Tdk210 and Tdk1 (27.7% identity)—showed no detectable drive (Supplementary Fig. 4). The robust drive activity of the Tdk210–Tdk11 chimeras indicates that their structure–function relationships have remained compatible over ∼100 million year of evolution. Together, these findings demonstrate that Tdk proteins share conserved structural features despite deep evolutionary divergence.

### *tdk* drivers share a post-germination killing mode that disrupts chromosome segregation

Unlike most characterized fungal KMDs, which act by killing spores^29^, *tdk1* exerts its killing effect after spore germination: toxic Tdk1 assemblies disrupt mitotic chromosome segregation in noncarrier progeny by generating aberrant chromosomal adhesions through Bdf1/Bdf2^21^. To determine whether *tdk210* and *tdk203* employ a similar post-germination killing mode, we performed live imaging of tetrads from *ade6::tdk210* × *ade6* and *ade6::tdk203* × *ade6* crosses. Like the situation with *tdk1*, all four spores from a tetrad appeared morphologically normal and underwent germination and germ tube elongation (Fig. 4a). However, the noncarrier spores from *ade6::tdk210* × *ade6* crosses typically arrested after the first mitotic division, whereas those from *ade6::tdk203* × *ade6* crosses often underwent several limited divisions before arresting (Fig. 4a,b).

**Fig. 4:**
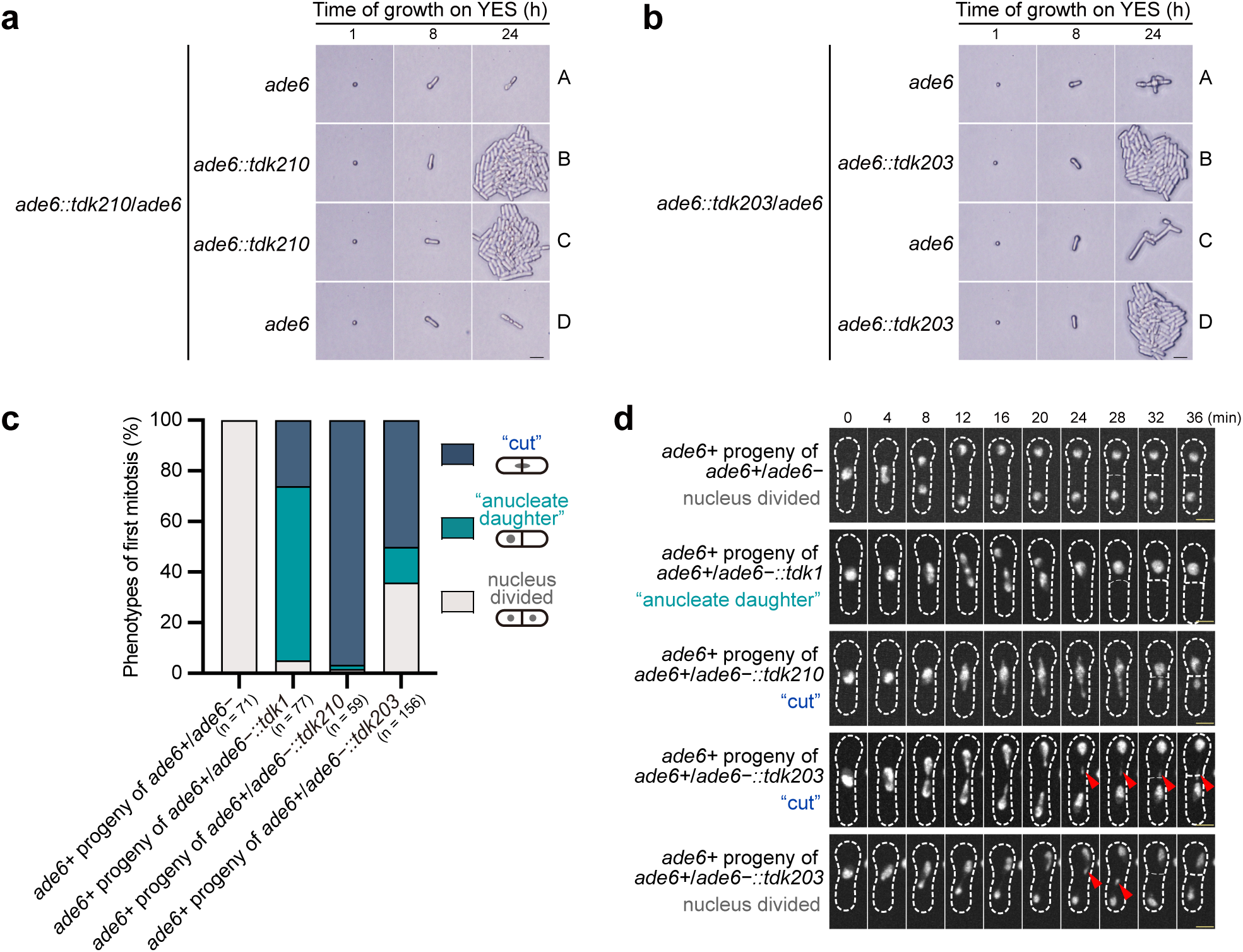
*tdk* drivers kill noncarrier progeny through post-germination disruption of chromosome segregation. **a,b**, Bright-field images of spores from *ade6::tdk210* × *ade6* (**a**) and *ade6::tdk203* × *ade6* (**b**) crosses; all four spores exhibit normal germination and germ tube elongation, but two noncarriers fail to proliferate. Genotypes were determined by replica-plating; the *tdk* insertion was linked to a *natMX* drug-resistance marker. A–D denote the four progeny from one ascus. Scale bar, 20 μm. **c,d**, Time-lapse fluorescence microscopy of chromosome segregation defects in *ade6+* progeny—derived from *ade6+* × *ade6−* (control) and *ade6+* × *ade6−::tdk* crosses—during the first post-germination mitosis. **c**, Quantification of mitotic cells displaying the indicated phenotypes: “cut”, “anucleate daughter”, and nucleus divided. Images were acquired every 15 min for quantification. n, number of mitotic cells analyzed. **d**, Representative time-lapse images of noncarriers. Notably, in *ade6+* × *ade6−::tdk203* crosses, even noncarrier progeny that completed nuclear division exhibited lagging chromosomes (red arrows) when imaged at higher temporal resolution (images acquired every 2 min, displayed at 4-min intervals). Chromatin was visualized using Bdf1-mECitrine. Scale bar, 2 μm.

We further performed time-lapse imaging of noncarrier progeny (*ade6+*, selectively germinated on adenine-deficient medium) from *ade6+* × *ade6−::tdk* crosses. All three *tdk* drivers caused chromosome segregation defects during the first post-germination mitosis, manifesting as either “cut” or “anucleate daughter” phenotypes (Fig. 4c,d). The distribution of these phenotypes varied among drivers: *tdk1*—68.8% anucleate daughter, 26.0% cut; *tdk210*—96.6% cut; *tdk203*—14.1% anucleate daughter, 50.0% cut. Notably, even those *tdk203* noncarrier cells that completed nuclear division exhibited lagging chromosomes when imaged at higher temporal resolution (Fig. 4d), indicating that toxic effects were exerted on chromosomes, albeit not strong enough to completely sabotage mitosis. Supporting these findings, vegetative cells expressing self-killing *tdk* mutants also displayed chromosome segregation defects (Supplementary Fig. 5). Together, these results demonstrate that *tdk* drivers share a conserved post-germination killing mode that disrupts mitotic chromosome segregation.

### *tdk1*, *tdk210*, and *tdk203* are mutually incompatible KMDs

We next tested whether *tdk* drivers confer cross-resistance to one another’s killing activity. In all pairwise crosses in a strain background lacking endogenous *tdk1*—*ade6::tdk1* × *ade6::tdk210*, *ade6::tdk1* × *ade6::tdk203*, and *ade6::tdk210* × *ade6::tdk203* — progeny inheriting either driver exhibited low viability (Fig. 5a). Consistent with this lack of cross-protection, both *tdk210* and *tdk203* retained full drive activity in a *tdk1+* background (Fig. 5b). When two *tdk* drivers were placed at unlinked loci, only double-carrier progeny survived robustly. In *tdk1+ ade6* × *tdk1Δ ade6::tdk210* crosses, double carriers (*tdk1+ ade6::tdk210*) accounted for 91% of viable progeny. Similarly, in *tdk1+ ade6* × *tdk1Δ ade6::tdk203* crosses, double carriers constituted 80% of viable progeny, and in *ura4 ade6::tdk203* × *ura4::tdk210 ade6* crosses, they made up 88% of viable progeny (Fig. 5c,d). These results demonstrate that *tdk1*, *tdk210*, and *tdk203* are mutually incompatible KMDs.

**Fig. 5:**
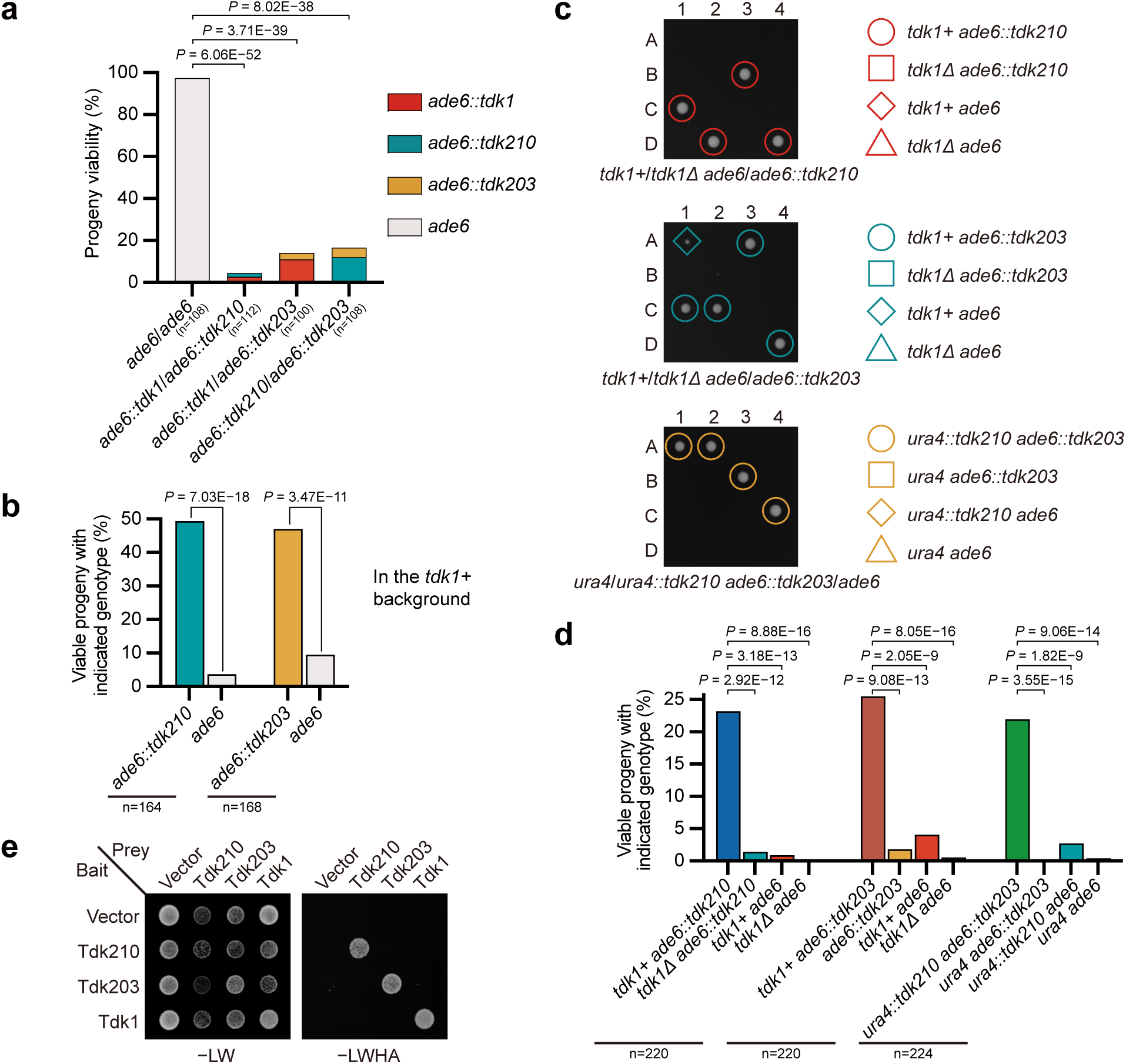
*tdk1*, *tdk210*, and *tdk203* are mutually incompatible KMDs. **a**, Tetrad analyses of *ade6* × *ade6* control crosses, *ade6::tdk1* × *ade6::tdk210*, *ade6::tdk1* × *ade6::tdk203*, and *ade6::tdk210* × *ade6::tdk203* crosses in a *tdk1Δ* background, showing no cross-resistance between different *tdk* drivers. **b**, Tetrad analyses of *ade6::tdk210* × *ade6* and *ade6::tdk203* × *ade6* crosses in a *tdk1+* background, demonstrating full drive activity of *tdk210* and *tdk203* in the presence of *tdk1*.**c,d**, Tetrad analyses of *tdk1+ ade6* × *tdk1Δ ade6::tdk210*, *tdk1+ ade6* × *tdk1Δ ade6::tdk203*, and *ura4 ade6::tdk203* × *ura4::tdk210 ade6* crosses, showing survival of only double carriers. **d**, Statistical analysis of progeny viability in **c**. **e**, Y2H assays demonstrating self-interaction of Tdk1, Tdk210, and Tdk203, but no detectable interaction between different Tdk proteins. *−*LW: SD/−Leu/−Trp; *−*LWHA: SD/−Leu/−Trp/−His/−Ade. For **a**, *P* values (Fisher’s exact test) compare progeny viability to the *ade6* × *ade6* control. For **b,d**, *P* values (exact binomial test) compare progeny viability of the two genotypes within each cross. n, total progeny analyzed.

We previously showed that, in its antidote conformation, Tdk1 colocalizes with and promotes the dissolution of supramolecular assemblies formed by Tdk1 in its toxin conformation^20^. Although the precise molecular mechanism remains unclear, a plausible hypothesis is that antidote function requires direct interaction with its cognate toxin, and that this specificity is enabled by the intrinsic self-interaction capability of Tdk1. Consistent with this model, yeast two-hybrid (Y2H) assays revealed robust homotypic interactions for Tdk1, Tdk210, and Tdk203, but no detectable heterotypic interactions among them (Fig. 5e). This link between specific physical interactions and resistance is further reinforced by the observation that Tdk11 confers resistance to Tdk210 and interacts with Tdk210 in Y2H assays (Supplementary Fig. 6). Collectively, these data indicate that *tdk1*, *tdk210*, and *tdk203* are mutually incompatible, likely because their antidotes cannot neutralize noncognate toxins due to a lack of cross-interaction.

### *tdk210*, *tdk203*, and *tdk11* execute killing independently of Bdf1/Bdf2

Although *tdk* drivers are functionally conserved and share overall structural similarity, sequence alignment revealed that the Bdf1/Bdf2-interacting region — essential for Tdk1-mediated killing—is poorly conserved in Tdk210, Tdk203, and Tdk11 (Fig. 6a). Consistent with this, Y2H assays showed that, unlike the C-terminal fragment of Tdk1 (Tdk1C), the corresponding fragments of Tdk210 (Tdk210C) and Tdk203 (Tdk203C) failed to interact with Bdf1 (Fig. 6b).

**Fig. 6:**
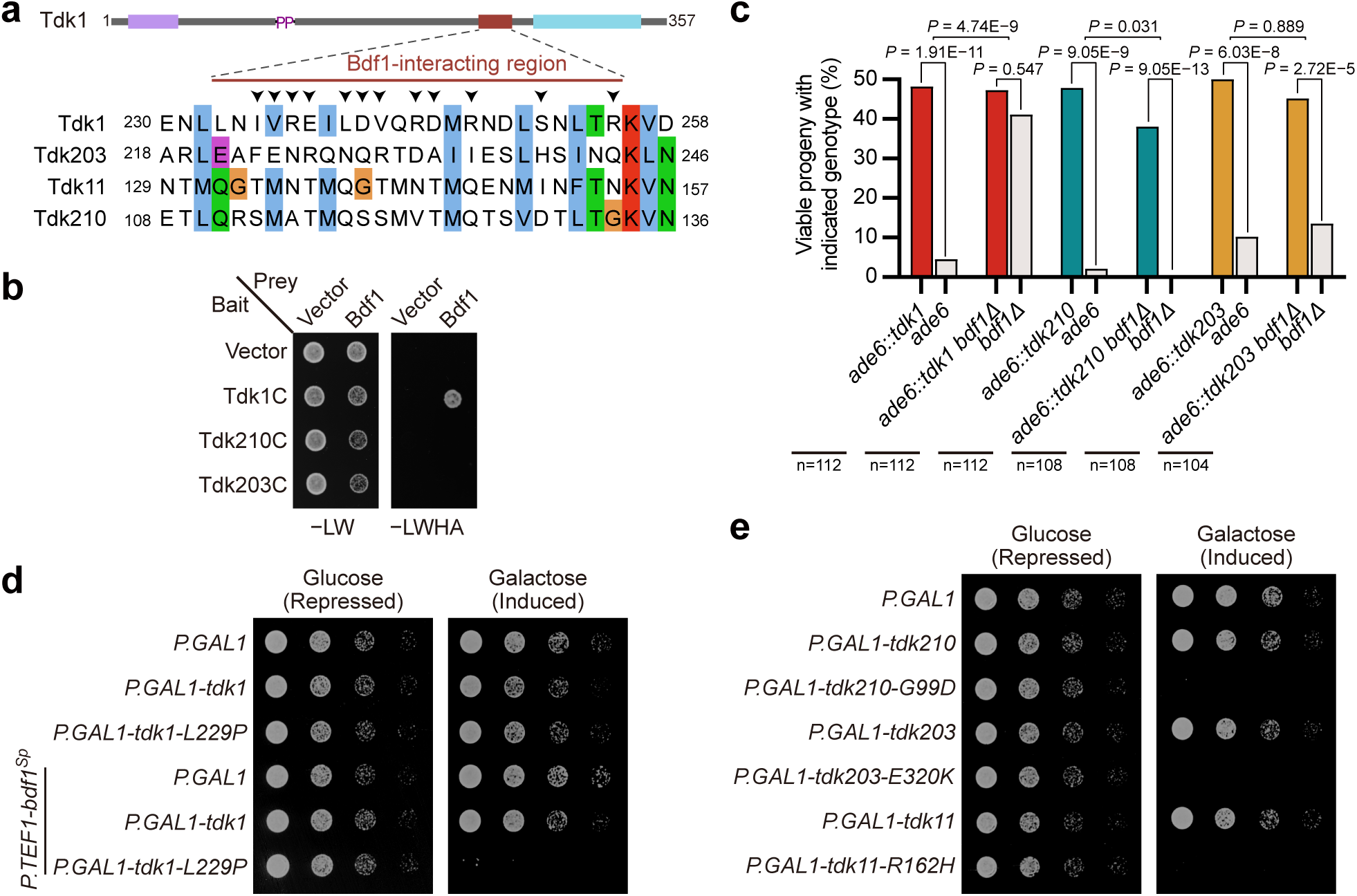
*tdk210*, *tdk203*, and *tdk11* mediate killing independently of Bdf1. **a**, Sequence alignment of the Bdf1-interacting region of Tdk1 with Tdk11, Tdk210, and Tdk203, showing poor conservsion of this region. Arrows indicate Tdk1 residues directly involved in Bdf1-binding^21^. **b**, Y2H assays showing interaction between Bdf1 and the C-terminal fragment of Tdk1 (Tdk1C, residues 227**–**357), but not with the corresponding fragments of Tdk210 (Tdk210C, 104**–**234) or Tdk203 (Tdk203C, 214**–**342). **c**, Tetrad analyses of *ade6::tdk1* × *ade6*, *ade6::tdk210* × *ade6*, and *ade6::tdk203* × *ade6* in *bdf1*+ and *bdf1Δ* backgrounds, demonstrating that killing by *tdk1*—but not by *tdk210* or *tdk203*—is reduced in the absence of *bdf1*. *P* values within individual crosses were calculated using the exact binomial test; *P* values between crosses were calculated using Fisher’s exact test. n, total progeny analyzed. **d**, Spot assays in *S. cerevisiae* showing that the toxicity of Tdk1-L229P (expressed form the galactose-inducible *P.GAL1* promoter) requires co-expression of *S. pombe* Bdf1 (Bdf1^Sp^), expressed from the constitutive *P.TEF1* promoter. **e**, Spot assays in *S. cerevisiae* showing that Tdk210-G99D, Tdk203-E320K, and Tdk11-R162H (but not their wild-type counterparts) are directly toxic when expressed from the *P.GAL1* promoter.

Tdk1-mediated progeny killing primarily depends on Bdf1, with Bdf2 playing a minor role in the absence of Bdf1^21^. To assess whether *tdk210* and *tdk203* requires Bdf1/Bdf2 for killing, we tested their drive efficiency in *bdf1Δ* and *bdf2Δ* backgrounds. Loss of *bdf1* dramatically suppressed *tdk1* killing, increasing noncarrier viability in *ade6::tdk1* × *ade6* crosses from 9.0% to 82.2% (Fig. 6c). In stark contrast, noncarrier viability remained low in both *ade6::tdk210* × *ade6* and *ade6::tdk203* × *ade6* crosses, with no significant difference between *bdf1+* and *bdf1Δ* backgrounds (Fig. 6c). Likewise, deletion of *bdf2* had no effect on the killing activity of *tdk210* or *tdk203* (Supplementary Fig. 7a).

Consistent with the essential role of Bdf1/Bdf2 in supporting Tdk1-mediated killing, heterologous expression of the self-killing Tdk1-L229P mutant in *Saccharomyces cerevisiae* showed no toxicity unless *S. pombe* Bdf1 was co-expressed (Fig. 6d), likely because the Tdk1-interacting region is poorly conserved in the *S. cerevisiae* homologs of Bdf1 and Bdf2 (Supplementary Fig. 7b). In contrast, the self-killing variants Tdk210-G99D, Tdk203-E320K, and Tdk11-R162H each caused robust growth inhibition in *S. cerevisiae* without requiring co-expression of any *S. pombe* cofactor (Fig. 6e), indicating that their host factors are conserved in this species.

Together, these findings demonstrate that—unlike *tdk1*—*tdk210*, *tdk203*, and *tdk11* execute killing independently of Bdf1/Bdf2.

### Evolutionary persistence and dynamics of the *tdk* family in *Schizosaccharomyces*

To trace the evolutionary trajectory of the *tdk* family in *Schizosaccharomyces*, we performed phylogenetic and synteny analyses. A maximum-likelihood tree of 44 intact *tdk* genes, based on the conserved IPR013902 domain, shows topological incongruence with the species tree (Fig. 7a), indicating a complex evolutionary history. The tree resolves four major clades (A–D): Clade A contains only *S. pombe tdk1* and its *S. pombe* CN ortholog *tdk110*; Clade B comprises many genes from non-*S. pombe* species; Clade C contains only three genes—*S. pombe tdk11* and its *S. pombe* CN ortholog *tdk109*, and *tdk210* from *S. cryophilus*; and Clade D includes many genes from both *S. pombe* and non-*S. pombe* species. Within Clades B and D, there are multiple occasions of closely related genes from the same species clustering together (e.g., *tdk404*, *tdk416*, and *tdk409* in Clade B and *tdk301* and *tdk303* in Clade D), indicating recent lineage-specific expansions—likely accounting for variable *tdk* gene numbers across taxa.

**Fig. 7:**
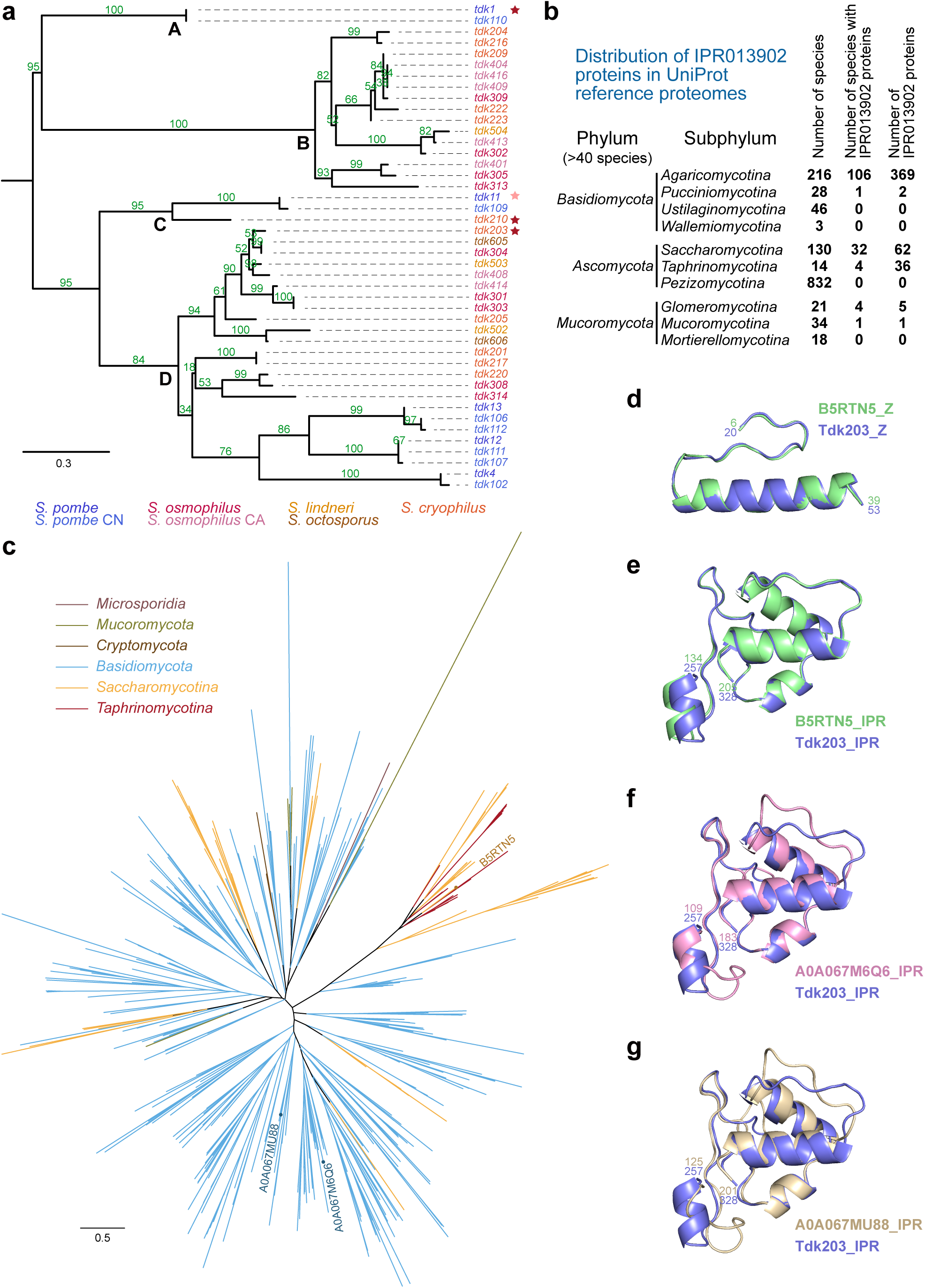
*tdk* genes constitute an ancient, highly diverged KMD family with deep evolutionary roots in fungi. **a**, Maximum-likelihood phylogenetic tree of 44 intact Tdk homologs from seven *Schizosaccharomyces* genomes, based on alignment of the IPR013902 domain (mid-point rooting). Four major clades (A–D) are labeled. Scale bar: 0.3 substitutions per site. Active drivers (*tdk1*, *tdk210*, *tdk203*) are marked with red stars; the marginal driver (*tdk11*) with a pink star. UFBoot)support values for each branch are indicated in green (%). **b**, Taxonomic distribution of IPR013902-containing proteins across fungal phyla with ≥ 40 reference species in UniProt reference proteomes (release 2025_04), displayed at the phylum and subphylum level. The full hierarchical distribution of 478 IPR013902-containing proteins across fungal species is provided in Supplementary Data 2. **c**, Maximum-likelihood phylogenetic tree of fungal IPR013902-containing proteins, inferred from IPR013902 domain alignment (mid-point rooting). Scale bar: 0.5 substitutions per site. Data are from UniProt reference proteomes (release 2025_04). The tree file is provided in Supplementary Data 3. **d–g**, Structural alignments of the Z-shaped domain (**d**) and IPR013902 domain (**e**) of B5RTN5 with Tdk203 (blue) (Z-domain RMSD = 0.402 Å over 188 atoms; IPR013902 domain RMSD = 0.337 Å over 382 atoms); and IPR013902 domain of A0A067M6Q6 (**f**; RMSD = 0.742 Å over 306 atoms) and A0A067MU88 (**g**; RMSD = 0.818 Å over 277 atoms) with Tdk203 (blue). Structures were predicted by AlphaFold 3^26^; full models and PAE plots are provided in Supplementary Fig. 10a–c.

Notably, *tdk304* (*S. osmophilus*) and *tdk605* (*S. octosporus*) — from species that diverged ∼30 Mya^22^—exhibited little sequence divergence (94.1% amino acid identity), suggesting potential horizontal gene transfer, a phenomenon previously documented for the *Spok* KMD family in filamentous fungi^15,16,30^. Active *tdk* drivers (*tdk1*, *tdk210*, and *tdk203*) are distributed across three of the four major clades, supporting that *tdk* genes in the common ancestor of these *Schizosaccharomyces* species—diverged 100 Mya—were already functional KMDs.

Synteny analysis revealed that cross-species synteny of *tdk* genes is rare. No syntenic *tdk* loci were detected between *S. pombe* and non-*S. pombe* species. Among non-*S. pombe* species, we identified five syntenic *tdk* sets. Two of these are conserved across the four members of the *S. osmophilus*–*S. lindneri*–*S. octosporus* clade (hereafter the *osmo–lind–octo* group)—*S. osmophilus*, *S. osmophilus* CA, *S. lindneri*, and *S. octosporus*—which share a common ancestor ∼16 Mya^22^: one set includes two intact genes and two pseudogenes (*tdk305*, *tdk401*, *tdk501\**, *tdk601\**); the other comprises four pseudogenes (*tdk310\**, *tdk405\**, *tdk505\**, *tdk602\**) (Supplementary Fig. 8a,b).

The remaining three syntenic sets extend to *S. cryophilus*, which diverged from the *osmo–lind–octo* group ∼30 Mya^22^, but exhibit lineage-specific gene loss. Two sets — *tdk211\**/*tdk314*/*tdk411\** and *tdk210*/*tdk315\**/*tdk410\** — are present in *S. cryophilus*, *S. osmophilus*, and *S. osmophilus* CA, but absent in *S. lindneri* and *S. octosporus* (Supplementary Fig. 8c,d). The third set, located near chromosome ends, shows local copy-number variation: *S. osmophilus* and *S. osmophilus* CA harbor tandem pseudogene pairs (*tdk311*:tdk312\** and *tdk406*:tdk407\**), whereas *S. cryophilus* carries a *wag* family gene—an uncharacterized gene family often found flanking *wtf* KMDs^17^—between two *tdk* genes (*tdk215\** and *tdk216*) (Supplementary Fig. 8e). In contrast, *S. octosporus* contains only a single pseudogene (*tdk603\**), and *S. lindneri* lacks any *tdk* gene at this locus (Supplementary Fig. 8e). These patterns indicate recurrent gene loss and occasional local duplication during *tdk* evolution.

To trace the degeneration trajectory of an active driver, we focused on the *tdk210*/*tdk315\**/*tdk410\** set, which includes the active driver *tdk210*. *tdk315\** in *S. osmophilus* isolates and *tdk410\** in *S. osmophilus* CA carry distinct frameshift and nonsense mutations (Supplementary Fig. 9a). Using *tdk210* and *tdk11* as outgroups, we reconstructed their intact ancestral sequence (*tdk315^ra^*; ra = reconstructed ancestor; see Methods). Despite strong *P.adh1*-driven expression, *tdk315^ra^* showed no drive activity but conferred substantial resistance to *tdk210*: *ade6::P.adh1-tdk315^ra^* progeny exhibited 75.6% viability (Supplementary Fig. 9b). This supports a model in which ancestral active *tdk* genes first lose toxin function while retaining antidote activity—a resistant allele—before accumulating degenerative mutations in natural populations^21^.

Consistent with irreversible toxin function loss, introducing the self-killing G99D mutation (at a conserved residue) into *tdk315^ra^*failed to restore toxicity, whereas the equivalent mutation rendered the marginal driver *tdk11* toxic (Supplementary Fig. 9c). Collectively, these data demonstrate the long-term persistence of the *tdk* KMD family in *Schizosaccharomyces*, accompanied by frequent functional decay, pseudogenization, and gene loss.

### *tdk* homologs exist in multiple fungal phyla

As noted earlier, the IPR013902 domain is the most conserved region in Tdk proteins. According to the InterPro database^31,32^, this domain is found exclusively in fungal proteins. To systematically explore the taxonomic distribution of IPR013902-containing proteins, we surveyed the UniProt reference fungal proteomes (1,430 fungal species)^33^ and identified a total of 478 IPR013902 proteins across 150 species—predominantly within the phyla *Basidiomycota* (371 proteins in 107 species), *Ascomycota* (98 proteins in 36 species), and *Mucoromycota* (6 proteins in 5 species) (Fig. 7b and Supplementary Data 2).

Within these phyla, the distribution of IPR013902 proteins is highly uneven and patchy. For example, among the 832 species in the *Ascomycota* subphylum *Pezizomycotina*, none contain IPR013902 proteins. In the two other *Ascomycota* subphyla, only 29% (4 out of 14) of *Taphrinomycotina* species—all of which are *Schizosaccharomyces* species—and 25% (32 out of 130) of *Saccharomycotina* species (none in *Saccharomycetaceae*) contain them. This highly discontinuous distribution suggests that IPR013902 proteins have been frequently lost during fungal evolution and/or acquired via horizontal gene transfer. Notably, species that do encode IPR013902 proteins often carry multiple copies; the highest number is found in the wood-decaying basidiomycete *Botryobasidium botryosum*, which encodes 17 IPR013902 proteins (Supplementary Data 2).

Phylogenetic analysis revealed that the *Taphrinomycotina* IPR013902 proteins (i.e., *Schizosaccharomyces* Tdk proteins) cluster with 18 proteins from the *Saccharomycotina* families *Debaryomycetaceae* and *Pichiaceae* (Fig. 7c and Supplementary Data 3), suggesting possible horizontal gene transfer between two *Ascomycota* subphyla. However, these sequences represent only a small fraction of the total diversity of the IPR013902 proteins across fungi (Fig. 7c). This pattern suggests that *Schizosaccharomyces* Tdk proteins descend from a profoundly ancient origin—likely predating the divergence of major fungal phyla.

Testing whether IPR013902 genes in non-*Schizosaccharomyces* species act as KMDs is challenging due to genetic intractability. We therefore selected two non-*Schizosaccharomyces* species—the *Saccharomycotina* species *Debaryomyces hansenii* and the aforementioned *Basidiomycota* species *B. botryosum* — and compared the genomic and structural features of their IPR013902 genes to those of the *Schizosaccharomyces tdk* family genes, which not only possess conserved functional and structural traits but also other salient signatures, including multiple genes per genome and high pseudogenization rates. *D. hansenii* encodes nine IPR013902 proteins, a substantial fraction of which lack a domain N-terminal to the IPR013902 domain—suggestive of pseudogenization. An apparently intact IPR013902 protein in this species, B5RTN5, exhibits high structural similarity to *Schizosaccharomyces* Tdk proteins in both the N-terminal Z-shaped domain and the IPR013902 domain (Fig. 7d,e and Supplementary Fig. 10a). Two of the 17 *B. botryosum* IPR013902 proteins (A0A067M6Q6 and A0A067MU88) possess an N-terminal region adopting a Z-shaped structure, an IPR013902 domain, and a PP motif in between, thus exhibiting structural resemblance to *Schizosaccharomyces* Tdk proteins (Fig. 7f,g and Supplementary Fig. 10b–e).

Notably, in the phylogenetic tree, IPR013902 proteins from both *D. hansenii* and *B. botryosum* are distributed across multiple, distantly related clades and exhibit greater sequence diversity than *Schizosaccharomyces* Tdk proteins (Supplementary Fig. 11)— a pattern suggestive of long-term evolutionary persistence and/or extensive horizontal acquisition. Collectively, the similarities in protein structure and genomic features suggest that many fungal IPR013902 genes in non-*Schizosaccharomyces* species may encode KMDs. Our findings are consistent with the emergence of an IPR013902-containing KMD in the last common ancestor of fungi, followed by extensive lineage-specific loss—a pattern mirroring the rapid turnover observed for *tdk* loci in *Schizosaccharomyces*. Thus, *tdk* family genes and their non-*Schizosaccharomyces* homologs may constitute an ancient and highly diverged KMD superfamily with an extraordinarily broad yet discontinuous phylogenetic distribution.

## DISCUSSION

Killer meiotic drivers (KMDs) have traditionally been viewed as evolutionarily ephemeral, restricted to narrow phylogenetic distributions^1,6,7,11,12^. Our study challenges this perspective by demonstrating that *Schizosaccharomyces tdk* genes represent an ancient and diverse KMD family. The identification and functional characterization of two active *tdk* drivers in *S. cryophilus*—*tdk210* and *tdk203*, which share only 27.7% and 27.3% amino acid identity, respectively, with the previously characterized *S. pombe* KMD *tdk1* — provide direct evidence for the long-term persistence of this KMD family. Despite extensive sequence divergence, these three active drivers share conserved structural features and a post-germination killing mode. Remarkably, despite sharing the trait of killing through disruption of mitotic chromosome segregation, they differ in their dependence on host factors. We propose that this diversification in host factor requirements, together with their mutual incompatibility, may reflect an evolutionary strategy that enables this KMD family to escape extinction through repeated cycles of functional innovation. In addition, we investigated the distribution and features of non-*Schizosaccharomyces tdk* homologs across multiple fungal phyla and identified characteristics supporting the possibility that some of them may also encode KMDs.

Mutual incompatibility among members of the same KMD family has been documented in other KMD families, such as *wtf* and *Spok*^16,18,34^. Our finding of mutual incompatibility among *tdk1*, *tdk210*, and *tdk203* suggests that, for single-gene KMDs requiring self-interaction for detoxification, evolutionary divergence can render family members unable to cross-interact, resulting in mutual incompatibility. This process parallels the evolution of oligomeric proteins, where gene divergence leads to new homomers that no longer interact with their progenitors^35,36^. The tempo of this incompatibility emergence remains an open question. On one hand, members that diverged ∼100 Mya still maintain interaction and resistance (e.g., *tdk11* and *tdk210*). On the other hand, incompatibility can arise readily: copy number expansion of the repeat in exon 6 of *wtf4* from *S. pombe var. kambucha* rendered it incompatible with the shorter-repeat *wtf4*^37^. Notably, *tdk* genes also harbor internal repeats. Determining whether and how repeat variation contributes to KMD incompatibility offers a compelling avenue to dissect how rapidly incompatibility can evolve.

KMDs often impose fitness costs on their hosts, selecting for host suppressors and fueling molecular arms races^1,6,7^. A striking finding is that *tdk210*, *tdk203*, and *tdk11* do not rely on the same host proteins as *tdk1* — potentially targeting distinct chromosomal factors — for their killing activity. Such differences in host factor utilization may explain the phenotypic variation among *tdk* drivers (Fig. 4c,d). While the selective advantage driving this divergence remains unclear, one possibility is that new members hijack alternative host factors to avoid competition with existing drivers or reduce fitness costs by circumventing overexploitation of a single host factor. An alternative explanation is that hosts may evolve suppressive mutations in the originally targeted host factor, prompting *tdk* drivers to evade suppression by evolving novel interaction interfaces with alternative host proteins. Identifying the specific host factors facilitating the drive activity of *tdk210* and *tdk203* will be crucial for dissecting the mechanisms underlying family divergence and advancing our understanding of KMD–host arms races.

Gene diversification is a fundamental source of genetic novelty and is increasingly recognized as a key mechanism underpinning KMD persistence^13,16,38–42^. Once a KMD becomes fixed in a population or is suppressed by host mechanisms, its transmission advantage is lost, leading to degeneration. We propose two diversification scenarios that can restore this advantage: (i) divergence into an incompatible KMD variant that escapes neutralization by a preexisting driver, thereby regaining transmission advantage; and (ii) evolution of novel interaction interfaces that evade host suppression—such as that caused by mutations disrupting KMD–host interactions—thereby restoring drive activity.

In summary, our findings highlight that KMDs are dynamic and significant components of eukaryotic genomes. Although individual KMD genes frequently degenerate, their families persist across vast evolutionary timescales by intermittently diversifying into novel variants that reclaim transmission advantage. This fuels recurrent “genomic wars,” profoundly shaping eukaryotic genome evolution. We anticipate that additional evolutionarily persistent KMD families await discovery.

## METHODS

### Strain and plasmid construction

Strains and plasmids used in this study are listed in Supplementary Data 4. *Schizosaccharomyces pombe* strains were constructed using standard protocols^43^. Genomic integrations at the *ade6* or *ura4* loci were carried out using plasmids derived from stable-integrating vectors (SIVs)^44^. We used an SIV plasmid targeting the *ura4* locus to tag Bdf1 with mECitrine inserted between amino acids 34 and 35, as described previously^21^. For heterologous expression of *tdk* genes under the control of the *tdk1* promoter (*P.tdk1*) and terminator (*T.tdk1*), a 1017-bp upstream region and a 387-bp downstream region of *tdk1* were used. For *tdk1* expression, a 337-bp region upstream region was used as previously described^21^. To assess *tdk210* and *tdk203* drive activity in their native genomic contexts, we cloned the following fragments: for *tdk210*, a 1000-bp upstream region, the coding sequence (CDS), and a 735-bp downstream region; for *tdk203*, a 1029-bp upstream region, the CDS, and a 922-bp downstream region.

For expression in *Saccharomyces cerevisiae*, *tdk* variants were placed under the control of the galactose-induced *P.GAL1* promoter and integrated at the *LEU2* locus using pRS305-derived plasmids. *S. pombe* Bdf1 (Bdf1^Sp^) was expressed from the constitutive *P.TEF1* promoter using a pSV606-derived plasmid integrated at the *URA3* locus.

### Sequence homology search and annotation of *tdk* genes

To identify *tdk* homologs across *Schizosaccharomyces* species, we performed PSI-BLAST^23^ searches (E-value threshold = 0.001; matrix = BLOSUM45; gap existence = 15; gap extension = 2) against the predicted proteins encoded by the nine recently assembled telomere-to-telomere genomes (Jia et al., manuscript in preparation), using the protein sequences of the three active drivers—*tdk1*, *tdk210*, and *tdk203*. Searches were repeated until convergence (no new hits with E-value ≤ 0.001). Candidate homologs were manually curated to distinguish intact genes from pseudogenes using Oxford Nanopore Technologies (ONT) transcriptome data (Jia et al., manuscript in preparation) and sequence alignments. A sequence was classified as a pseudogene if it met either of the following criteria: (i) it contained inactivating mutations (frameshift or nonsense mutations); or (ii) it exhibited terminal truncation, defined by homologous flanking regions (>50 bp) identified in tBLASTn searches. Sequences lacking these features were classified as intact genes.

To identify additional *tdk* sequences, we performed tBLASTn searches (E-value threshold 1e−5) using the 44 intact Tdk protein sequences as queries against all nine genomes. Overlapping or contiguous hits were merged into single loci, and merged sequences shorter than 200 bp were excluded. For pseudogenes, genomic coordinates were manually refined based on tBLASTn hits. Detailed information for each *tdk* locus—including systematic IDs, gene symbols, and genomic coordinates—is provided in Supplementary Data 1.

### Synteny analysis

To identify cross-species synteny sets of *tdk* loci among *Schizosaccharomyces* species (*S. pombe*, *S. pombe* CN, *S. cryophilus*, *S. osmophilus*, *S. osmophilus* CA, *S. lindneri*, and *S. octosporus*), we used the *Schizosaccharomyces* Orthogroup (SOG) resource (https://fsnibs10.github.io/SOG/) to inspect syntenic blocks anchored by each *tdk* locus via its immediate upstream and downstream genes. If an immediate flanking gene lacked a conserved ortholog in other species, we iteratively examined the next adjacent gene outward until an orthologous anchor was identified or until the chromosomal end was reached. Synteny plots were generated using Clinker (https://cagecat.bioinformatics.nl/tools/clinker)^45^.

### De novo protein structure prediction and analysis

Protein structures were predicted using AlphaFold 3^26^ (seed = 1) via the AlphaFold server (https://alphafoldserver.com). For each target, the top-ranked model among the five generated models was used for subsequent structural analysis and visualization. Structural superposition and graphics were generated in PyMOL^46^, with visual properties adjusted to highlight key features. Root-mean-square deviation (RMSD) values for selected pairwise alignments were computed using PyMOL’s *align* command.

Structure-based sequence alignment (Fig. 6a and Supplementary Fig. 3e,f) of the four active or marginally active Tdk driver proteins (Tdk1, Tdk210, Tdk203, and Tdk11) were generated using FoldMason^47^.

### Taxonomic distribution of IPR013902 domain-containing proteins

Custom Python scripts were employed to query the UniProt REST API (UniProt release 2025_04) to enumerate, for each of the 1,430 fungal UniProtKB reference proteomes (proteome_type:1, taxonomy_id:4751), the number of proteins annotated with the InterPro entry IPR013902. All queries were executed on October 18, 2025. The results were used to generate a hierarchical fungal taxonomy tree (Supplementary Data 2), displaying the number of IPR013902 proteins for each species. The scripts and their outputs are available at https://github.com/lilindu/cryo-tdk-killers.

### Ancestral sequence reconstruction of *tdk315^ra^*

The putative ancestral coding sequence *tdk315^ra^* was inferred using a parsimony-based approach. Nucleotide sequences included three representative *S. osmophilus* strains and the *S. osmophilus* CA lineage. *tdk210* and *tdk212\** from *S. cryophilus* were used as outgroups to mitigate confounding effects arising from widespread inactivating mutations within *S. osmophilus*. Each nucleotide position was reconstructed independently as follows: (I) If a single nucleotide state was conserved across all *S. osmophilus* sequences, that state was assigned as ancestral. (II) If multiple states were present in *S. osmophilus*, the *S. cryophilus* outgroup sequences were used to infer the ancestral state: (i) If both *S. cryophilus* sequences shared an identical nucleotide, this consensus was assigned as ancestral—even if it differed from all *S. osmophilus* alleles; (ii) If the two *S. cryophilus* sequences differed, the ancestral state was inferred by allele matching with *S. osmophilus* (when each *S. cryophilus* allele matched one of the two *S. osmophilus* alleles, the allele corresponding to *tdk210* was designated ancestral; when only one *S. cryophilus* allele matched an *S. osmophilus* allele, that conserved allele was assigned as ancestral); (iii) For positions beyond 539 bp — corresponding to the nonconserved C-terminal region of *tdk212\**)—*tdk212\** was excluded from the analysis and the *tdk210* nucleotide state was assigned as ancestral if it did not match any *S. osmophilus* allele. The resulting ancestral sequence represents the most parsimonious reconstruction at each site. Predicted frameshift and nonsense mutations are indicated in Supplementary Fig. 9a.

### Sequence alignments and phylogenetic tree construction

For phylogenetic analysis of proteins harboring the IPR013902 domain in fungal proteomes, we refined the dataset (478) by replacing an initial set of 36 Tdk sequences from four *Schizosaccharomyces* species with a curated subset of 27 intact Tdk members obtained in this study. Amino acid sequence alignments for Fig. 7a,c and Supplementary Fig. 7b were generated using the E-INS-i algorithm implemented in the MAFFT v7 web server (https://mafft.cbrc.jp/alignment/server/)^48^. For phylogenetic analyses in Fig. 7a,c and Supplementary Fig. 11, sequences corresponding to the conserved IPR013902 domain were extracted using Jalview 2.11^49^. Maximum-likelihood phylogenetic trees were inferred with IQ-TREE v3.0^50^. Branch support values shown in Fig. 7a represent Ultrafast Bootstrap (UFBoot) support (%) based on 1,000 replicates. Among ten independent IQ-TREE runs, the best tree was selected for visualization in FigTree v1.4.4 (https://tree.bio.ed.ac.uk/software/figtree/).

For Supplementary Fig. 9a, coding sequences were aligned using MUSCLE, followed by manual refinement to maintain codon alignment integrity. A maximum-likelihood phylogenetic tree was inferred from this curated alignment using IQ-TREE and rooted by midpoint rooting. The final cladogram was visualized using FigTree.

### Tetrad analysis

Progeny viability was assessed by tetrad dissection using a TDM50 dissection microscope. Haploid parental strains of opposite mating types, freshly cultured on YES agar plates, were mixed on SPA agar plates and incubated at 30°C for 25–35 h to induce mating and sporulation. Mature asci were transferred to YES plates for microdissection. Dissected tetrads were incubated at 30°C for 3 days, and colony formation was documented by plate scanning. Genotypes were determined by replica plating.

To induce *tdk* expression during germination via the *P.tetO7* promoter in Fig. 2b, tetrads were digested onto YES plates supplemented with 5 μg/mL anhydrotetracycline (ahTet; Sigma, Cat#37919). Quantification of tetrad analysis results is presented in Supplementary Data 5.

### Spot assays

Log-phase cultures were adjusted to an OD_600_ of 0.4 for the initial spot, followed by fivefold serial dilutions. Cell suspensions were spotted onto solid media using a 48-pin frogger. Plates were incubated at 30°C and scanned after 2 days.

For toxicity assays in *S. pombe*, Tdk variants were expressed from the thiamine-repressible *P.41nmt1* promoter^51^. Cells were pre-grown in EMM medium supplemented with 5 μg/mL thiamine (Sinopharm, Cat#67002134), washed twice with sterile water, serially diluted, and spotted onto EMM plates either with (to repress expression) or without thiamine (to induce expression).

For toxicity assays in *S. cerevisiae*, Tdk variants were expressed under the *P.GAL1* promoter^52^. Log-phase cells pre-grown in sucrose-based SD medium were washed twice with sterile water, serially diluted, and spotted onto SD plates containing either 2% glucose (repression) or 2% galactose (induction).

### RCA mutagenesis

Random mutagenesis of the *tdk* genes was performed using error-prone rolling circle amplification (RCA)^27^. Reactions contained 1 ng of pDUAL-P.41nmt1-tdk plasmid template, phi29 DNA Polymerase (NEB, Cat#M0269L), and 2 mM MnCl₂ to promote replication errors. Following incubation at 30°C for 24 h, RCA products were digested with DpnI (to degrade methylated template plasmid DNA) and NotI (to generate ends homologous to the *leu1* genomic locus for targeted integration). The digested DNA were transformed into a *leu1-32 tdk1Δ tdk11Δ* strain and selected on leucine-deficient medium with thiamine (5 μg/mL). Resulting colonies were replica-plated onto thiamine-free media to induce *tdk* expression. Mutant toxicity was assessed by Phloxine B staining (5 μg/mL); enhanced red pigmentation indicated cellular toxicity. Clones exhibiting pronounced toxicity on thiamine-free plates were isolated for sequencing. The *tdk* coding region from selected clones was PCR-amplified and subjected to Sanger sequencing to identify causative mutations.

### Yeast two-hybrid (Y2H) assay

Y2H assays were performed using the Matchmaker Gold Yeast Two-Hybrid System (Clontech). Transformants co-expressing bait and prey constructs were selected on SD/−Leu/−Trp (−LW) media. Selected colonies were suspended in sterile water, adjusted to an OD_600_ of 1.0, and spotted onto SD/−Leu/−Trp/−His/−Ade (−LWHA) plates to assess activation of the *HIS3* and *ADE2* reporter genes. Plates were incubated at 30°C for 2–4 days before scanning.

### Fluorescence microscopy

Imaging was performed using an Andor Dragonfly 200 spinning-disk confocal system mounted on a Leica DMi8 inverted microscope, equipped with a 100×/1.40 NA oil-immersion objective. Acquired images were processed and analyzed using Fiji^24^.

For time-lapse experiments, cells were suspended in adenine-deficient EMM medium to selectively allow germination of *ade6*+ progeny, immobilized on microscope slides via agar pads (adenine-deficient EMM), and imaged at 30°C.

For imaging shown in Supplementary Fig. 5, cells were pre-incubated in EMM supplemented with 5 μg/mL thiamine, washed twice with sterile water, and then shifted to thiamine-free medium for 14–22 h at 30°C prior to imaging. Nuclei were stained using SYTOX Green nucleic acid stain (1 μM; Invitrogen, Cat#S7020) following fixation in 70% ethanol.

### Statistical analysis

Statistical analyses of tetrad quantification data are reported in Supplementary Data 6. Statistical data were visualized using GraphPad Prism, and exact *P* values are reported throughout. Detailed descriptions of statistical tests are provided in the corresponding figure legends. Deviations from the expected 1:1 Mendelian segregation ratio were evaluated using exact binomial tests (implemented in an Excel spreadsheet from BioStat Handbook: http://www.biostathandbook.com/exactgof.html)^53^. Comparisons of progeny viability and transmission ratio distortion between crosses were performed using Fisher’s exact tests ( http://www.biostathandbook.com/fishers.html)^53^.

## SUPPLEMENTAL INFORMATION

**Supplementary Figures 1–11.**

**Supplementary Data 1**: Genomic locus information of *tdk* genes in seven *Schizosaccharomyces* species.

**Supplementary Data 2**: Taxonomic distribution of IPR013902 domain-containing proteins across fungal reference proteomes.

**Supplementary Data 3**: Phylogenetic tree file for IPR013902 domain-containing proteins in fungal reference proteomes.

**Supplementary Data 4**: Strains and plasmids used in this study.

**Supplementary Data 5**: Quantification of tetrad analysis results.

**Supplementary Data 6**: Statistical analysis of tetrad quantification data.

## ACKNOWLEDGEMENTS

We thank Chen-Yu Fan, Yue Liang, Yu-Sheng Yang, Guang-Can Shao, and Yan-Hui Xu for technical assistance, and Yan Ding, Zhao-Qian Pan, Shao-Kai Ning, and Man-Yun Yang for sharing reagents. This work was supported by intramural funding from the National Institute of Biological Sciences and the Tsinghua Institute of Multidisciplinary Biomedical Research, Tsinghua University (L.-L. D.) and by the National Natural Science Foundation of China grant 31900405 (Y.H.). The funders had no roles in study design, data collection and analysis, decision to publish, or preparation of the manuscript.

## AUTHOR CONTRIBUTIONS

Conceptualization: F.-Y.Z., L.-L.D., Y.H.; Methodology and investigation: F.-Y.Z., G.-S.J., J.-Y.R., F.S., T.-Y.D., W.-C.Z., W.L., Q.-M.W., L.-L.D., Y.H. Writing – original draft: F.-Y.Z., L.-L.D., Y.H.; Writing – review and editing: F.-Y.Z., L.-L.D., Y.H.; Funding acquisition: L.-L.D. and Y.H.

---

**Supplementary Fig. 1:**
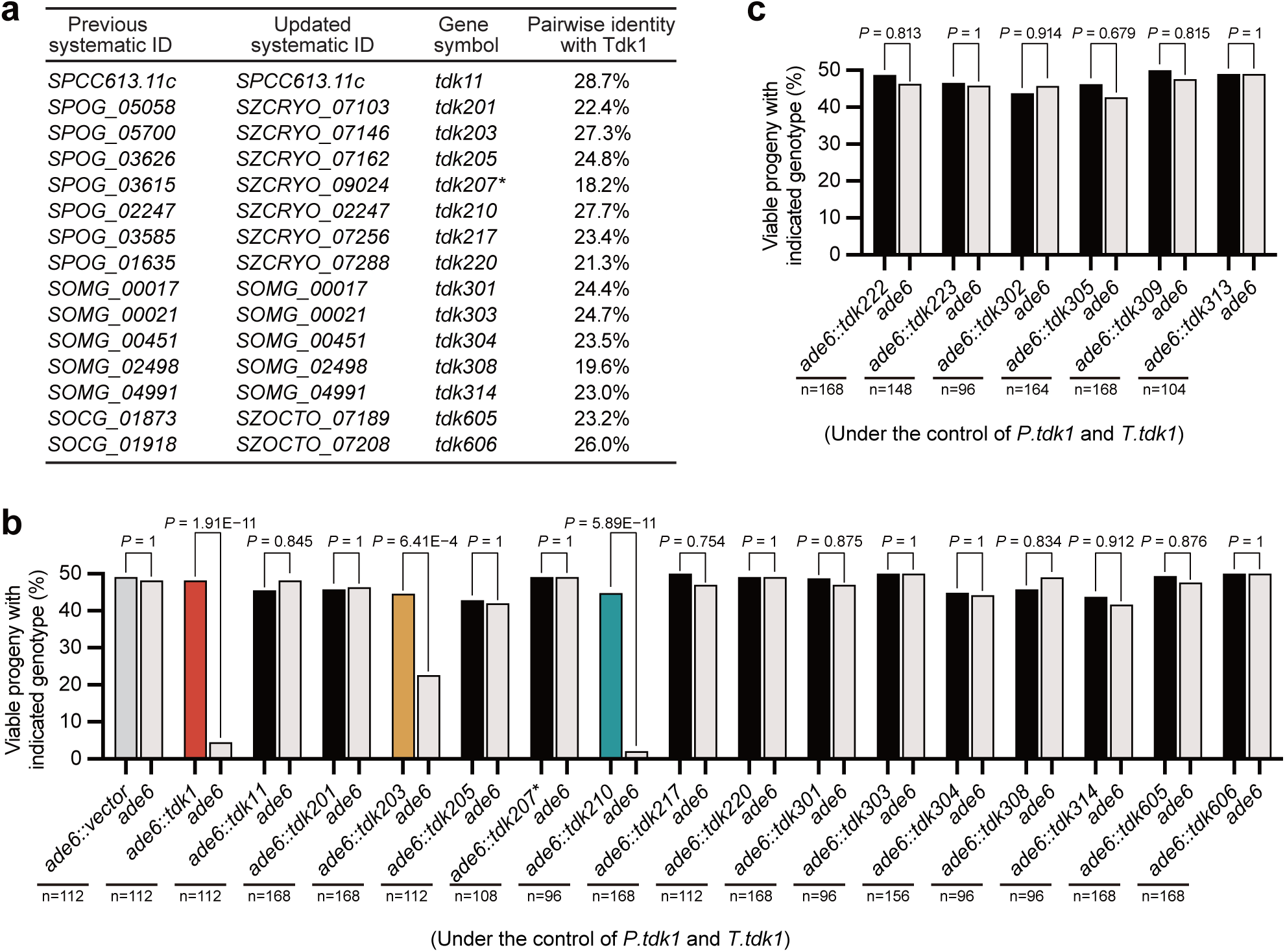
Identification of active KMDs among *tdk* homologs across fission yeasts. **a**, Pairwise amino acid identity of 15 homologs with Tdk1 (calculated using EMBL-EBI Needle^54^), with previous RefSeq systematic ID, updated systematic ID, and new gene symbols. **b,c** Tetrad analyses of 15 previously identified *tdk* homologs (**b**) and six additional *tdk* genes from *S. cryophilus* and *S. osmophilus* (**c**), revealing *tdk210* and *tdk203*—and no other homologs—as active KMDs. Each gene was controlled by *tdk1* promoter (*P.tdk1*) and terminator (*T.tdk1*), and integrated at the *ade6* locus in a *tdk1Δ tdk11Δ* background. *P* values (exact binomial test) compare observed viable progeny counts of the two genotypes to the expected 1:1 Mendelian segregation ratio. n, total progeny analyzed.

**Supplementary Fig. 2:**
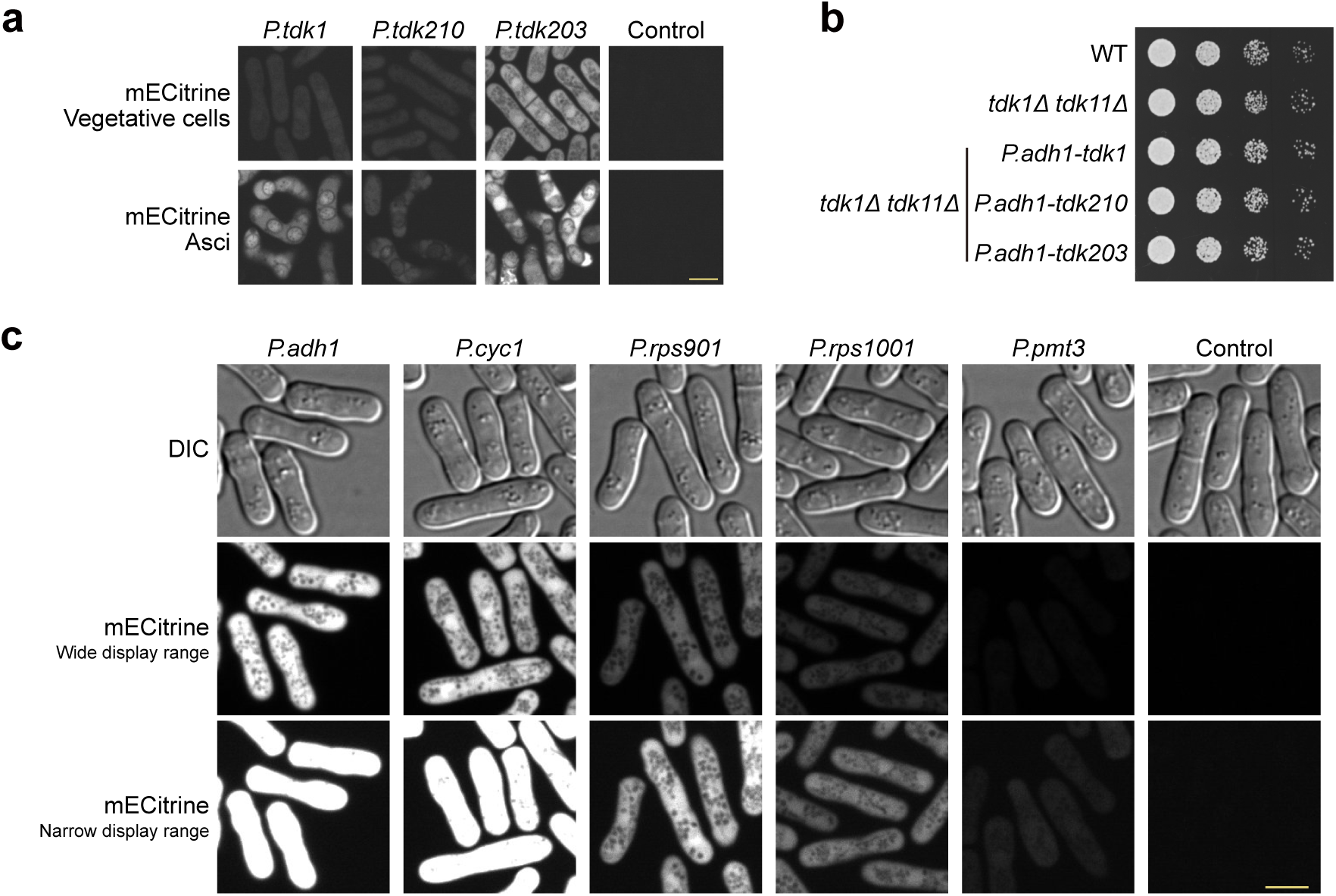
Transcriptional regulation of *tdk* drivers. **a**, Fluorescence micrographs of vegetative cells and asci, showing meiotic upregulation of mECitrine expression driven by *P.tdk1* but not by *P.tdk210* or *P.tdk203*. Strains carried the indicated *ade6*-integrated *promoter-mECitrine* constructs in a *tdk1Δ tdk11Δ* background. Asci were obtained by crossing each strain with a *tdk1Δ tdk11Δ* strain. Scale bar, 5 μm. **b**, Spot assays showing no vegetative toxicity of *tdk1*, *tdk210*, or *tdk203* expressed from the strong constitutive promoter *P.adh1* on the rich medium (YES). **c**, Fluorescence micrographs illustrating the relative expression strengths of the constitutive promoters *P.adh1*, *P.cyc1*, *P.rps901*, *P.rps1001*, and *P.pmt3* used in Fig. 2c. Scale bar, 5 μm.

**Supplementary Fig. 3:**
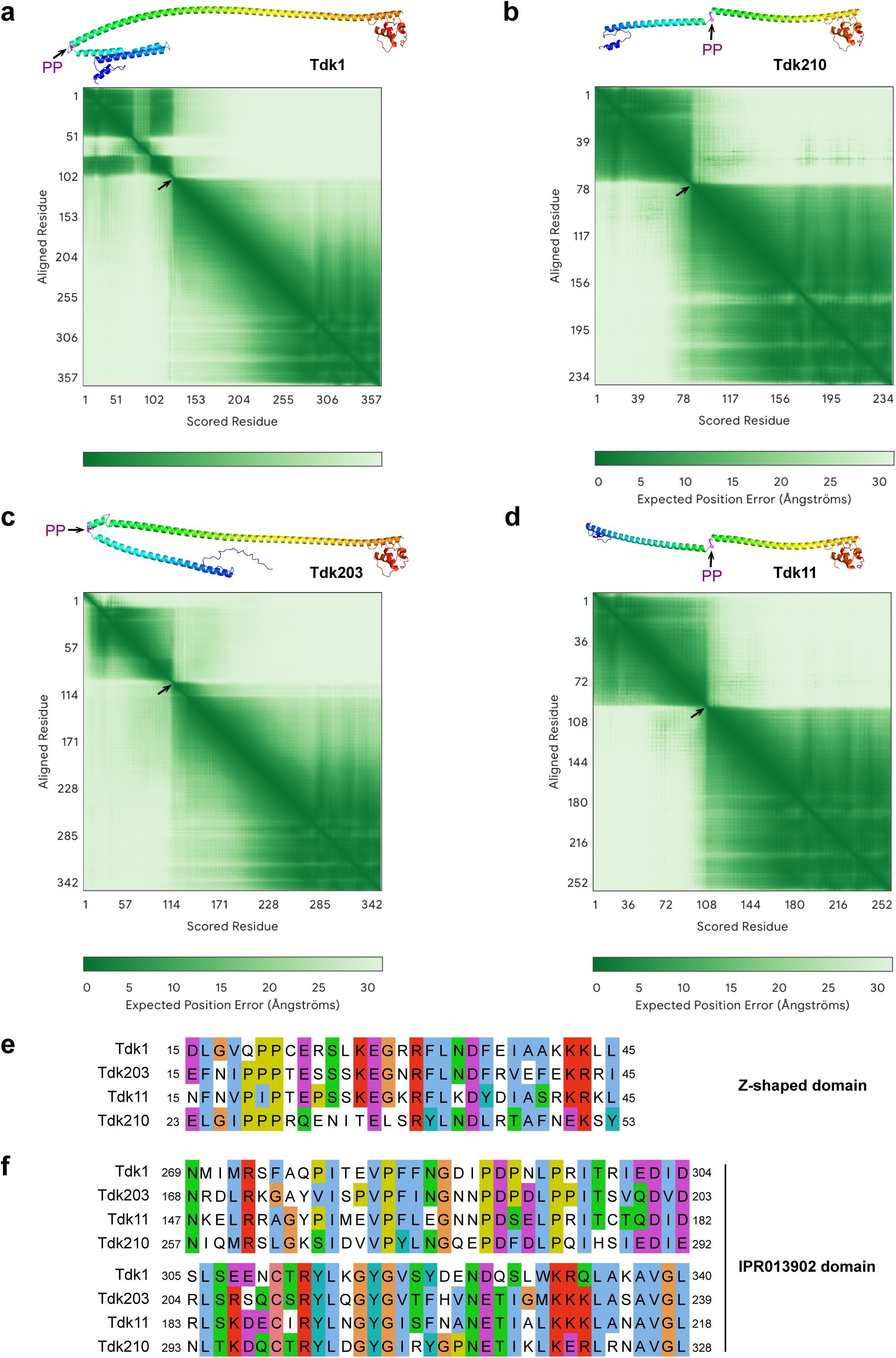
Structural predictions and conservation of Tdk driver proteins. **a–d**, AlphaFold 3^26^-predicted monomeric structures of Tdk1 (**a**), Tdk210 (**b**), Tdk203 (**c**), and Tdk11 (**d**), with corresponding predicted aligned error (PAE) plots shown on the right. Structures are rainbow-colored from N- to C-terminus; the conserved PP motif is highlighted in magenta (stick representation) and indicated by a black arrow in the PAE plots. **e,f** Sequence alignment of the conserved regions among Tdk1, Tdk203, Tdk210, and Tdk11: the N-terminal Z-shaped domain (**e**) and the IPR013902 domain (**f**).

**Supplementary Fig. 4:**
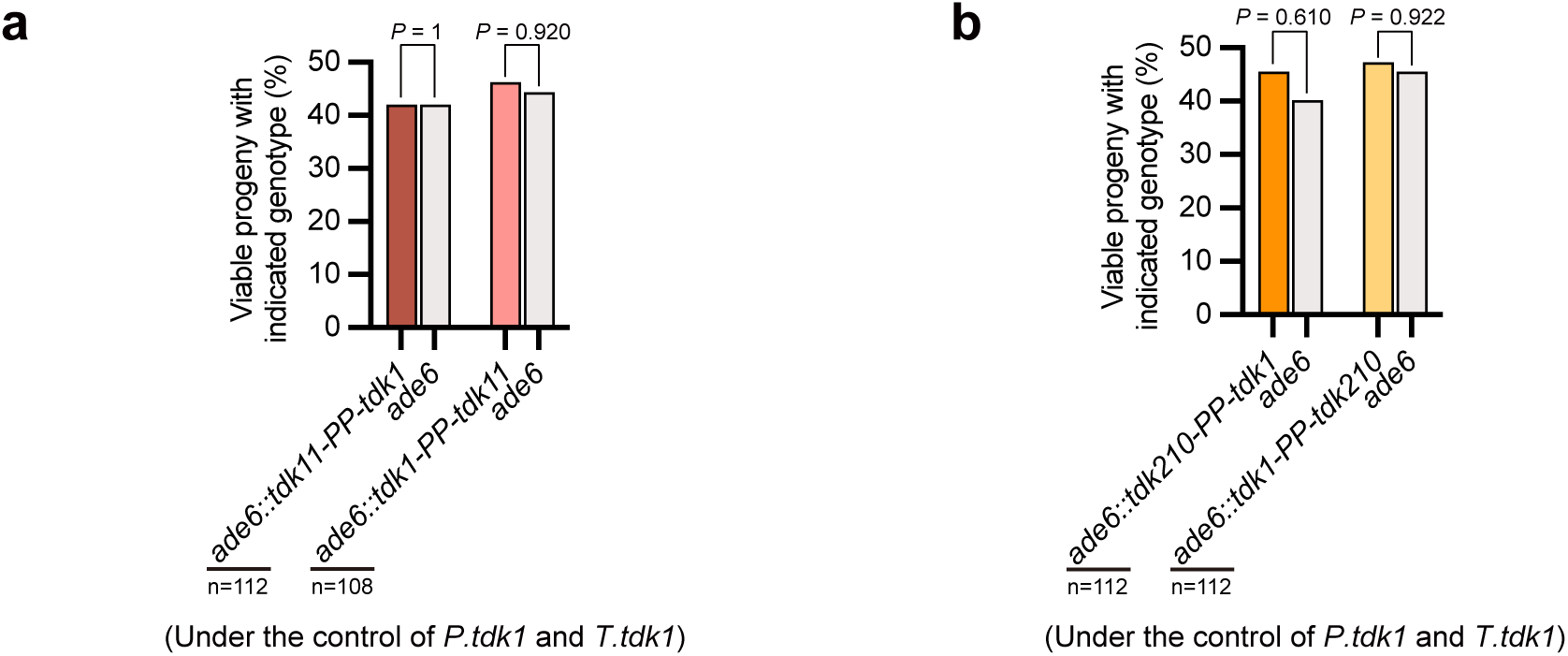
*tdk11*–*tdk1* and *tdk210*–*tdk1* chimeras lack drive activity. **a,b** Tetrad analyses of *tdk11***–***tdk1* (**a**) and *tdk210***–***tdk1* (**b**) chimeras (*tdk11-PP-tdk1*, *tdk1-PP-tdk11*, *tdk210-PP-tdk1*, *tdk1-PP-tdk210*) showing no drive activity. All constructs were under the control of *P.tdk1* and *P.tdk1*. *P* values (exact binomial test) compare progeny viability of the two genotypes within each cross. n, total progeny analyzed.

**Supplementary Fig. 5:**
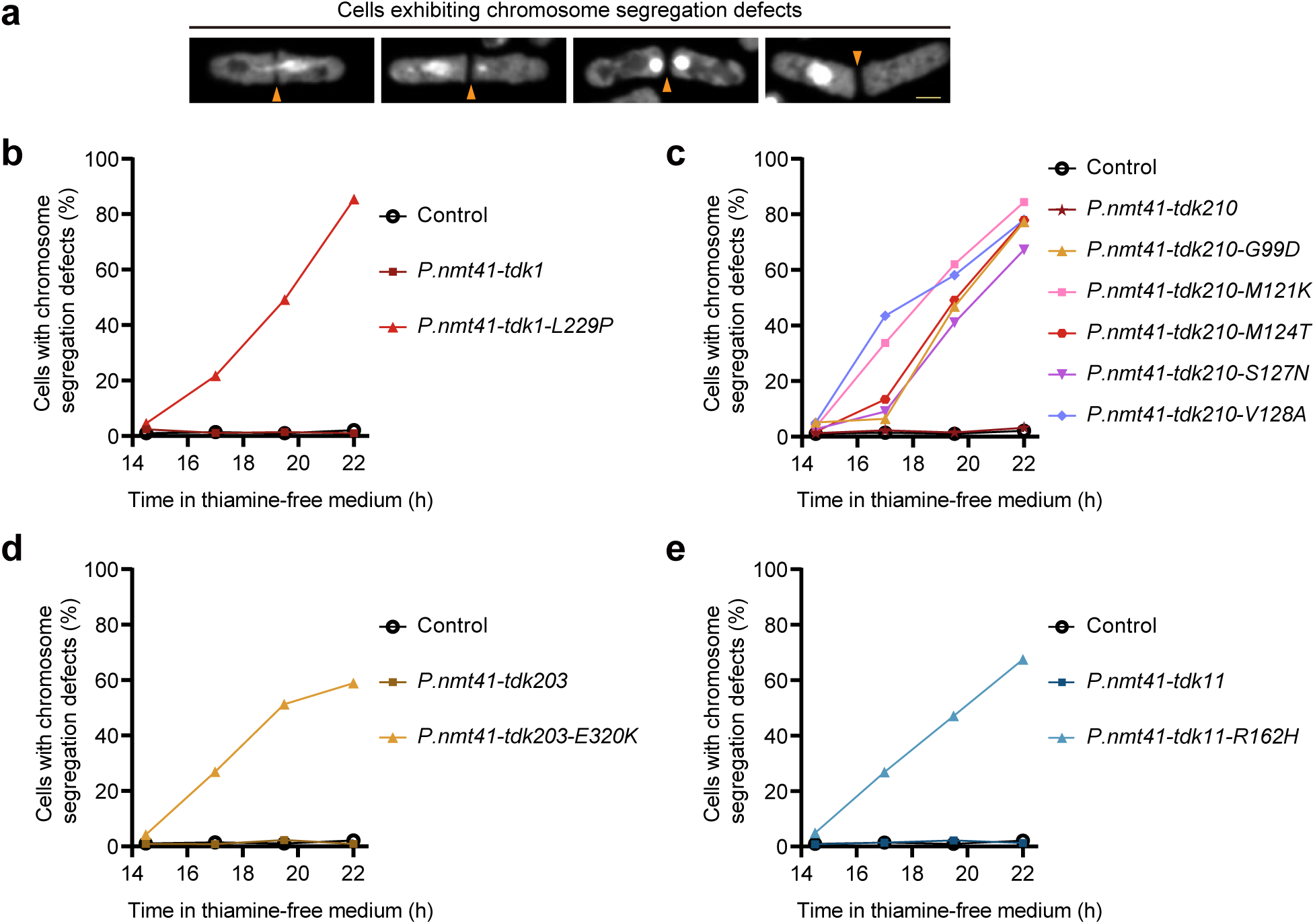
Self-killing *tdk* mutants disrupt mitotic chromosome segregation. **a**, Representative micrographs of cells with chromosome segregation defects upon expression of self-killing Tdk variants. Nuclei were stained with SYTOX Green nucleic acid stain; orange arrows indicate septa. Scale bar, 2 μm. **b–e**, Quantification of cells exhibiting chromosome segregation defects at the indicated time points following thiamine removal, which induces expression from the thiamine-repressible *P.nmt41* promoter^51^. Shown are wild-type and self-killing *tdk* variants whose expression was induced: *tdk1* and *tdk1-L229P* (**b**); *tdk210* and *tdk210-G99D*/*M121K*/*M124T*/*S127N*/*V128A* (**c**); *tdk203* and *tdk203-E320K* (**d**); *tdk11* and *tdk11-R162H* (**e**).

**Supplementary Fig. 6:**
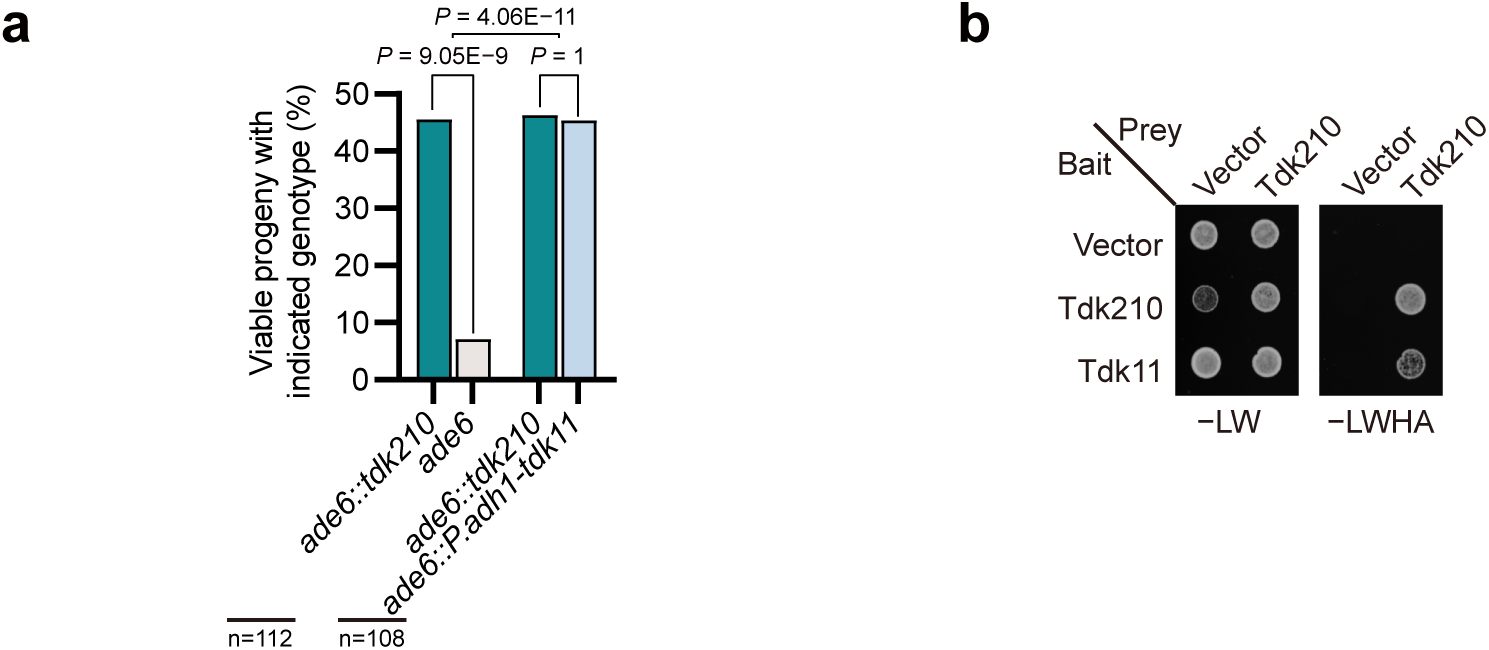
Tdk11 confers resistance to *tdk210*-mediated killing and interacts with Tdk210. **a**, Tetrad analyses of *ade6::P.adh1-tdk11* × *ade6::tdk210* and *ade6* × *ade6::tdk210* (control) crosses, showing full resistance of *P.adh1-tdk11* to *tdk210-*mediated killing. *P* values within individual crosses were calculated using the exact binomial test; The *P* value between crosses were calculated using Fisher’s exact test. n, total progeny analyzed. **b**, Y2H assays demonstrating a physical interaction between Tdk11 and Tdk210.

**Supplementary Fig. 7:**
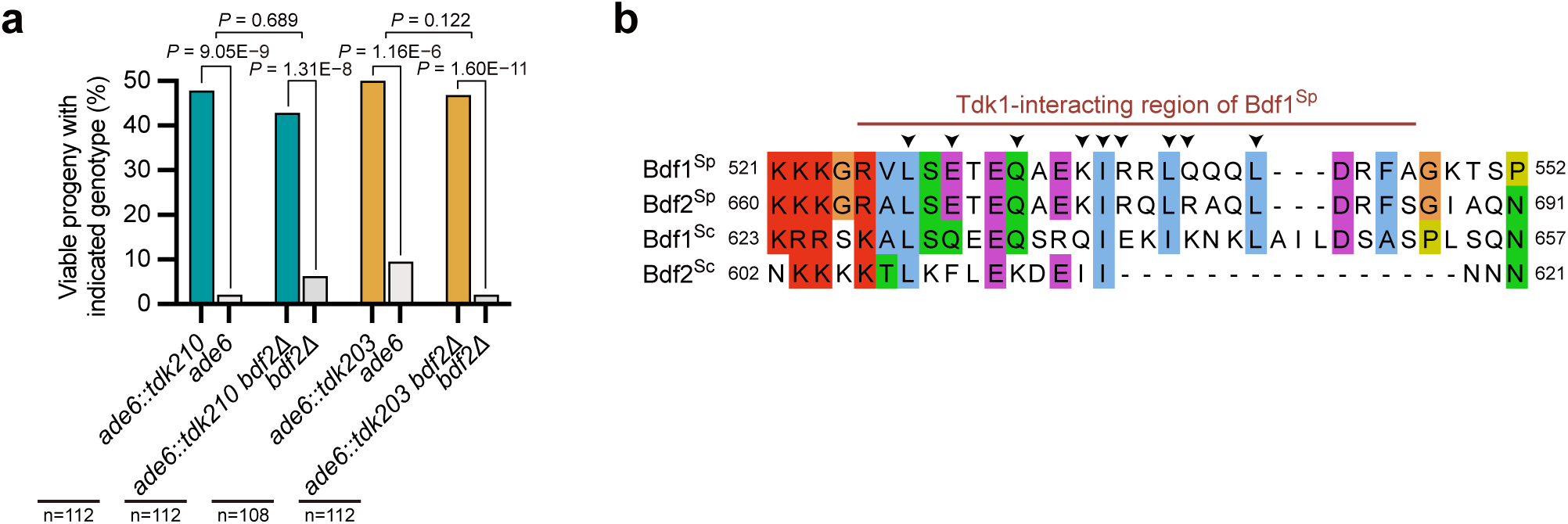
*tdk210* and *tdk203* drive independently of Bdf2. **a**, Tetrad analyses of *ade6::tdk210* × *ade6* and *ade6::tdk203* × *ade6* in *bdf2+* and *bdf2Δ* backgrounds, showing full drive activity of *tdk210* and *tdk203* in the absence of *bdf2*. *P* values within individual crosses were calculated using the exact binomial test; *P* values between crosses were calculated using Fisher’s exact test. n, total progeny analyzed. **b**, Sequence alignment of the Tdk1-binding regions of *S. cerevisiae* Bdf1 (Bdf1^Sc^) and Bdf2 (Bdf2^Sc^) with their orthologs in *S. pombe* (Bdf2^Sp^ and Bdf1^Sp^), showing that the corresponding regions in Bdf1^Sc^ and Bdf2^Sc^ exhibit low sequence conservation compared to those in *S. pombe*. Arrows indicate residues in Bdf1^Sp^ that directly interact with Tdk1^21^.

**Supplementary Fig. 8:**
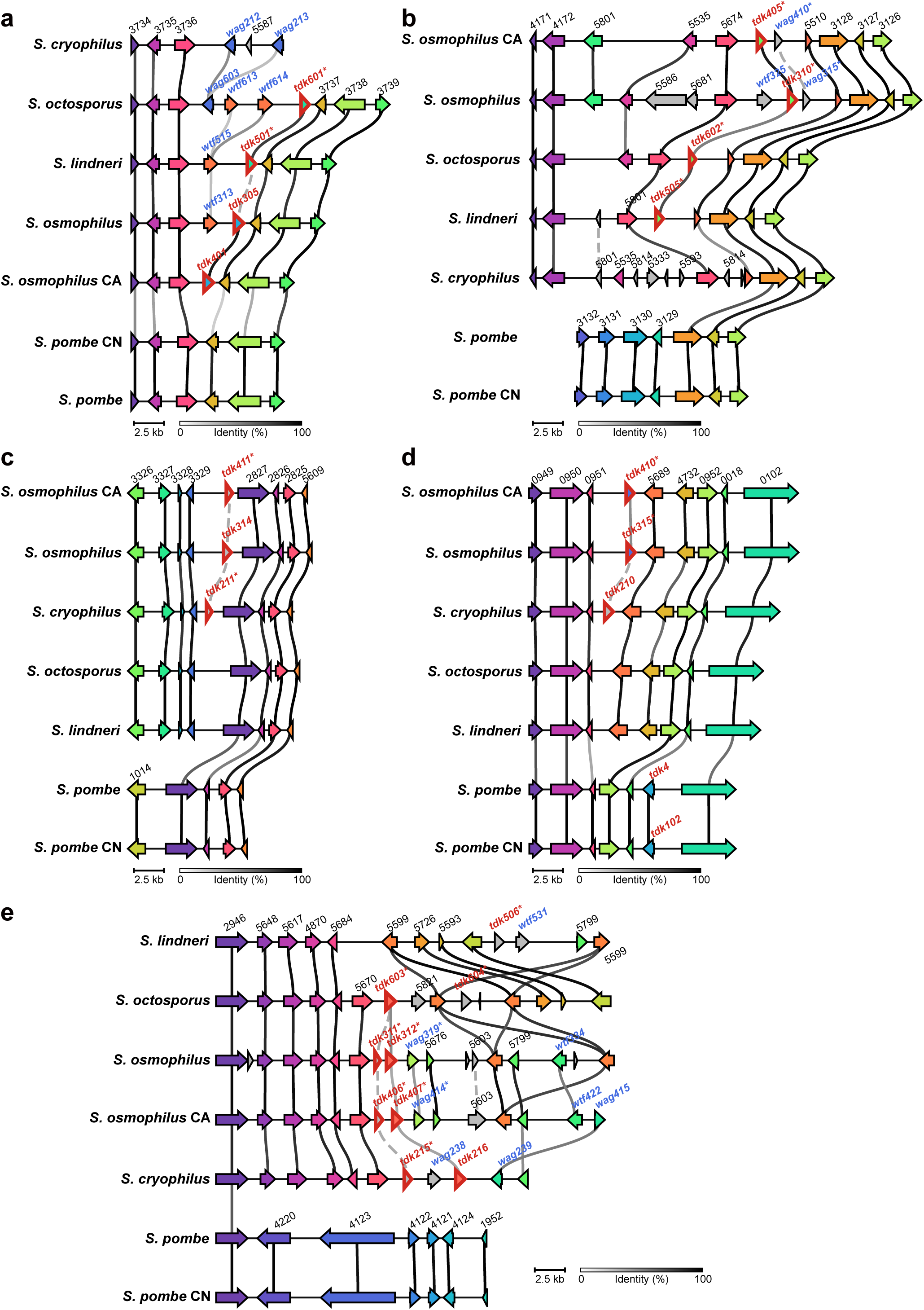
Cross-species synteny of *tdk* gene loci in the *Schizosaccharomyces* genus. **a–e**, Synteny analysis identified five cross-species synteny sets of *tdk* loci: two within the *osmo–lind–octo* group (*S. osmophilus*, *S. osmophilus* CA, *S. lindneri*, and *S. octosporus*) (**a,b**); two shared among *S. cryophilus*, *S. osmophilus*, and *S. osmophilus* CA (**c,d**); and one conserved in *S. cryophilus*, *S. osmophilus*, *S. osmophilus* CA, and *S. octosporus* (**e**). Four-digit numbers denote synteny group IDs from the *Schizosaccharomyces* Orthogroup (SOG) resource (https://fsnibs10.github.io/SOG/). *wtf* and *wag* genes, frequently found flanking *tdk* loci, are also labeled. Synteny plots were generated using Clinker^45^. Conserved genomic blocks are connected by solid lines, with line color indicating sequence identity. Only homologous regions sharing ≥ 30% sequence identity are linked by solid lines; connections below this threshold were manually added as gray dashed lines.

**Supplementary Fig. 9:**
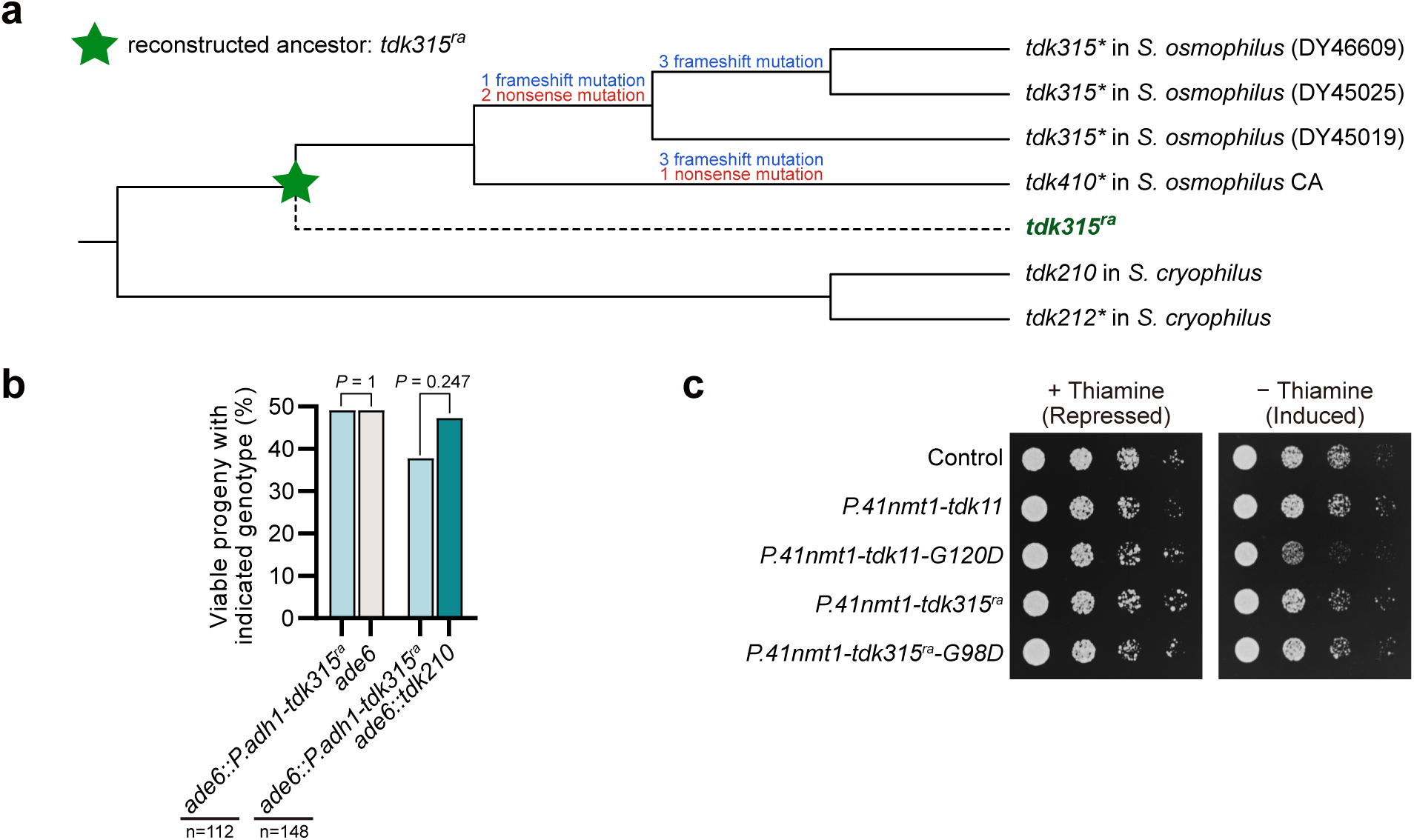
The ancestral state of *tdk315* is likely a resistant allele. **a**, Schematic of ancestral sequence reconstruction of *tdk315^ra^*. *S. osmophilus* CA and natural isolates of *S. osmophilus* carry distinct inactivating mutations (frameshift and nonsense mutations), consistent with degenerative evolution. *tdk210* and *tdk212\** from *S. cryophilus* were used as outgroups for ancestral inference (see Methods). The reconstructed ancestral sequence is designated *tdk315^ra^*. **b**, Tetrad analyses of *ade6::P.adh1-tdk315^ra^* × *ade6* and *ade6::P.adh1-tdk315^ra^* × *ade6::tdk210* crosses, showing that *P.adh1-tdk315^ra^* lacks drive activity but confers strong resistance to *tdk210-*mediated killing. *P* values (exact binomial test) compare progeny viability of the two genotypes within each cross. n, total progeny analyzed. **c**, Spot assays showing toxicity of *tdk11-G120D* but not *tdk315^ra^-G98D* when expressed from the *P.nmt41* promoter. The G120D substitution in *tdk11* and G98D in *tdk315^ra^* correspond to the self-killing G99D mutation in *tdk210*.

**Supplementary Fig. 10:**
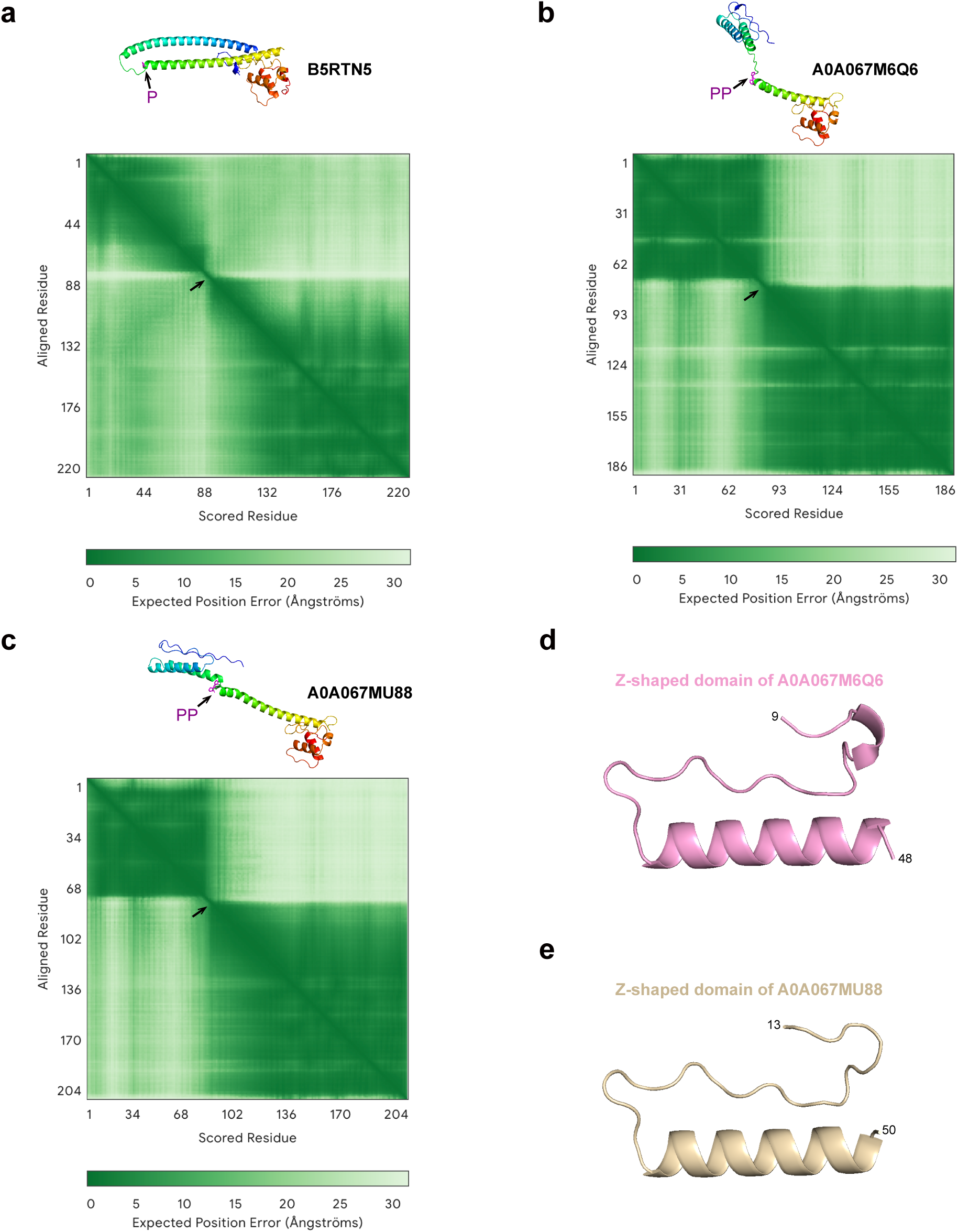
Structural predictions and similarity of non-*Schizosaccharomyces* Tdk homologs to *Schizosaccharomyces* Tdk driver proteins. a–c, AlphaFold 3^26^-predicted monomeric structures of B5RTN5 (a), A0A067M6Q6 (b), and A0A067MU88 (c), with PAE plots shown on the right. Structures are rainbow-colored from N- to C-terminus; the conserved PP motif is highlighted in magenta (stick representation) and indicated by a black arrow in the PAE plots. B5RTN5 contains a single proline at the position corresponding to the PP motif of *Schizosaccharomyces* Tdk driver proteins. **d,e**, Predicted N-terminal Z-shaped structure of A0A067M6Q6 (**d**) and A0A067MU88 (**e**), similar to that of *Schizosaccharomyces* Tdk driver proteins, despite the failure of structural superposition.

**Supplementary Fig. 11:**
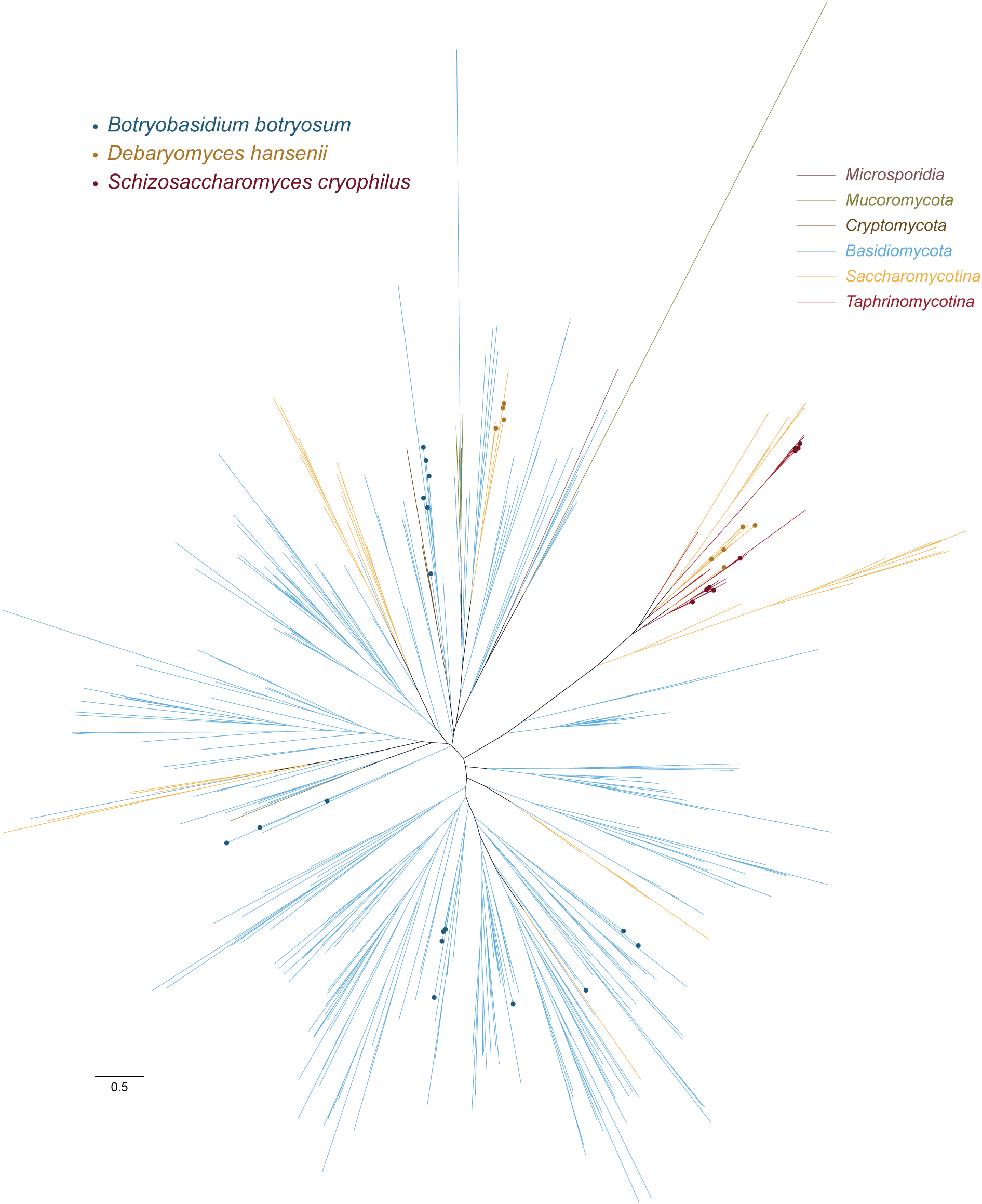
*tdk* homologs in *Debaryomyces hansenii* and *Botryobasidium botryosum* exhibit greater sequence divergence than those in *Schizosaccharomyces cryophilus*. Maximum-likelihood phylogenetic tree of fungal IPR013902-containing proteins (Fig. 7c), highlighting 9 *D. hansenii*, 17 *B. botryosum*, and 11 *S. cryophilus* homologs. The tree file is provided in Supplementary Data 3. Scale bar: 0.5 substitutions per site.

